## Supplementary Info for "Anatomy-to-Tract Mapping Infers White Matter Pathways Without Diffusion Streamline Propagation"

**Supplementary Note: Architectural Details of ATM**

The segmentation module of ATM comprises a total of $C$ 3D convolutional downsampling blocks, followed by $C-1$ upsampling blocks. The entire module is complemented by a long-skip connection for precise bundle segmentation. These convolutional blocks employ a kernel size of 3, and each block is equipped with filters specified as {32, 64, 128, 256}, followed by upsampling blocks with filter sizes of {128, 64, 32}. The final upsampling block is followed by a 3D convolutional output layer with a filter size of 1. The encoder-decoder component of the streamline VAE module is structured with $C$ 1D convolutional downsampling and upsampling blocks. This module utilizes a kernel size of 63, a latent space dimensionality of 64, and filter size for each downsampling and upsampling block are set as {32, 64, 128, 256} and {256, 128, 64, 32}, respectively. Each upsampling block is coupled with a FiLM layer[41] for adaptive feature modulation except the output layer with a filter size of 3. These FiLM layers receive input from the FiLM shared-weight block, which consists of two layers of multi-layer perceptron (MLP).

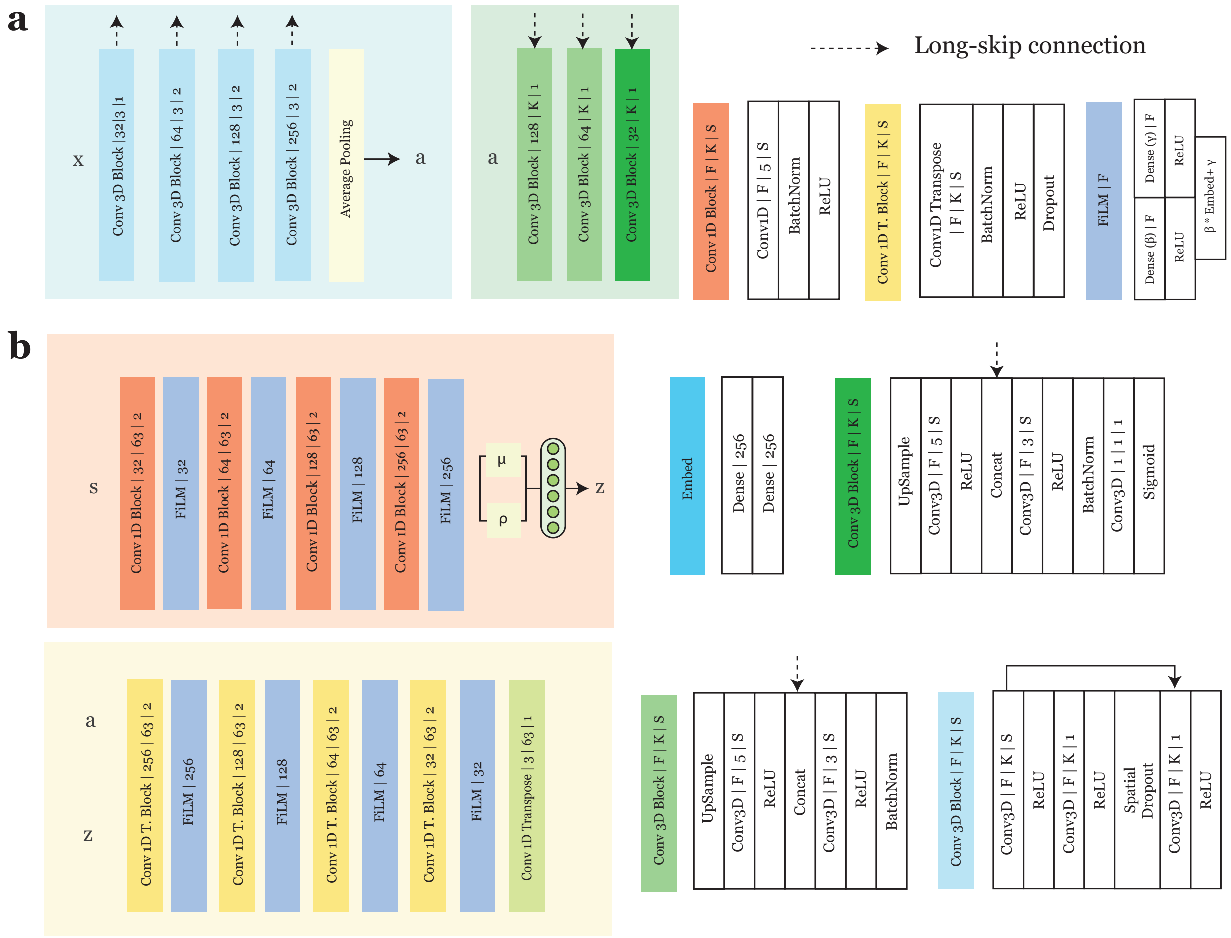


**Fig. S1** | **ATM architectural details**. **a**, Network architecture of T1w encoder, $E_A$, and decoder, $D_A$. **b**, Network architectures of streamline encoder, $E_S$, and decoder, $D_S$.

**Tab. S1** | White matter bundles.

| **Bundle** | **Abbreviation** |
|---|---|
| Arcuate fascicle | AF |
| Corpus callosum, Frontal lobe (most anterior part) | CC_Fr_1 |
| Corpus callosum, Frontal lobe (most anterior part) | CC_Fr_2 |
| Corpus callosum, Occipital lobe | CC_Oc |
| Corpus callosum, Parietal lobe | CC_Pa |
| Corpus callosum, Pre/Post central gyri | CC_Pr_Po |
| Cingulum | CG |
| Frontal aslant tract | FAT |
| Fronto-pontine tract | FPT |
| Inferior fronto-occipital fasciculus | IFOF |
| Inferior longitudinal fasciculus | ILF |
| Middle cerebellar peduncle | MCP |
| Middle longitudinal fascicle | MdLF |
| Optic radiation and Meyer's loop | OR_ML |
| Parieto-occipito pontine tract | POPT |
| Pyramidal tract | PYT |
| Superior longitudinal fasciculus | SLF |
| Uncinate fasciculus | UF |

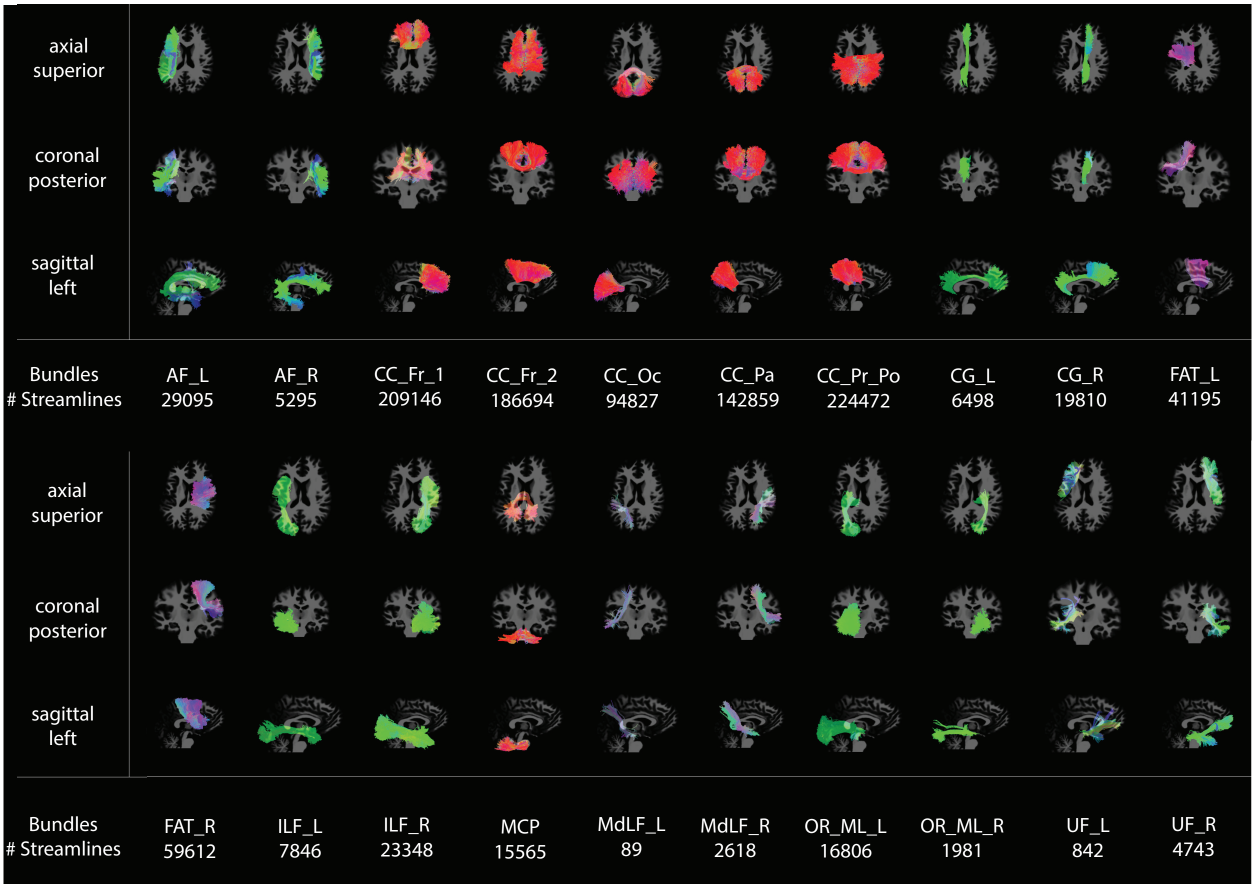


**Fig. S2 | Bundle comparison.** Ground truth bundles from Tractoinferno for a single representative subject.

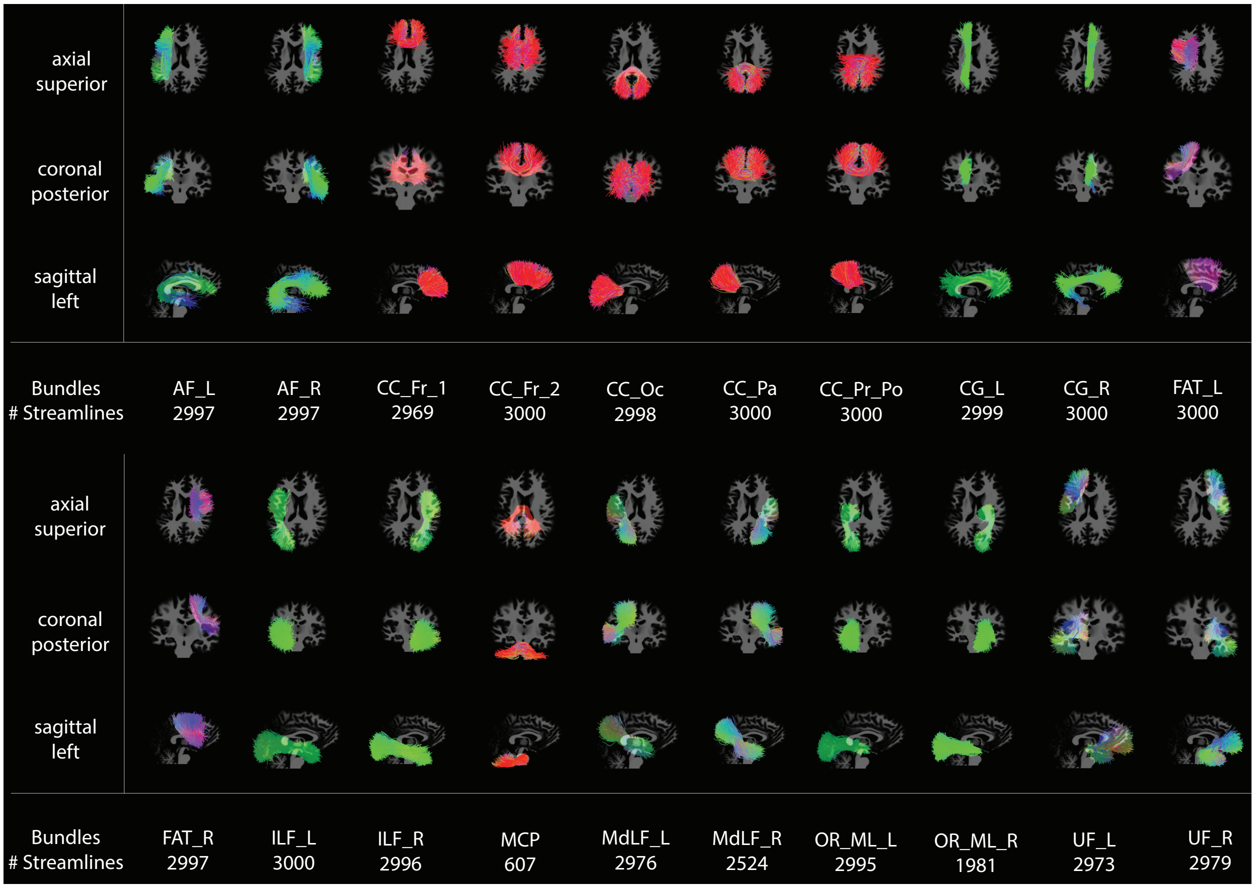


**Fig. S3 | Bundle comparison.** Bundles generated with ATM for a single representative subject.

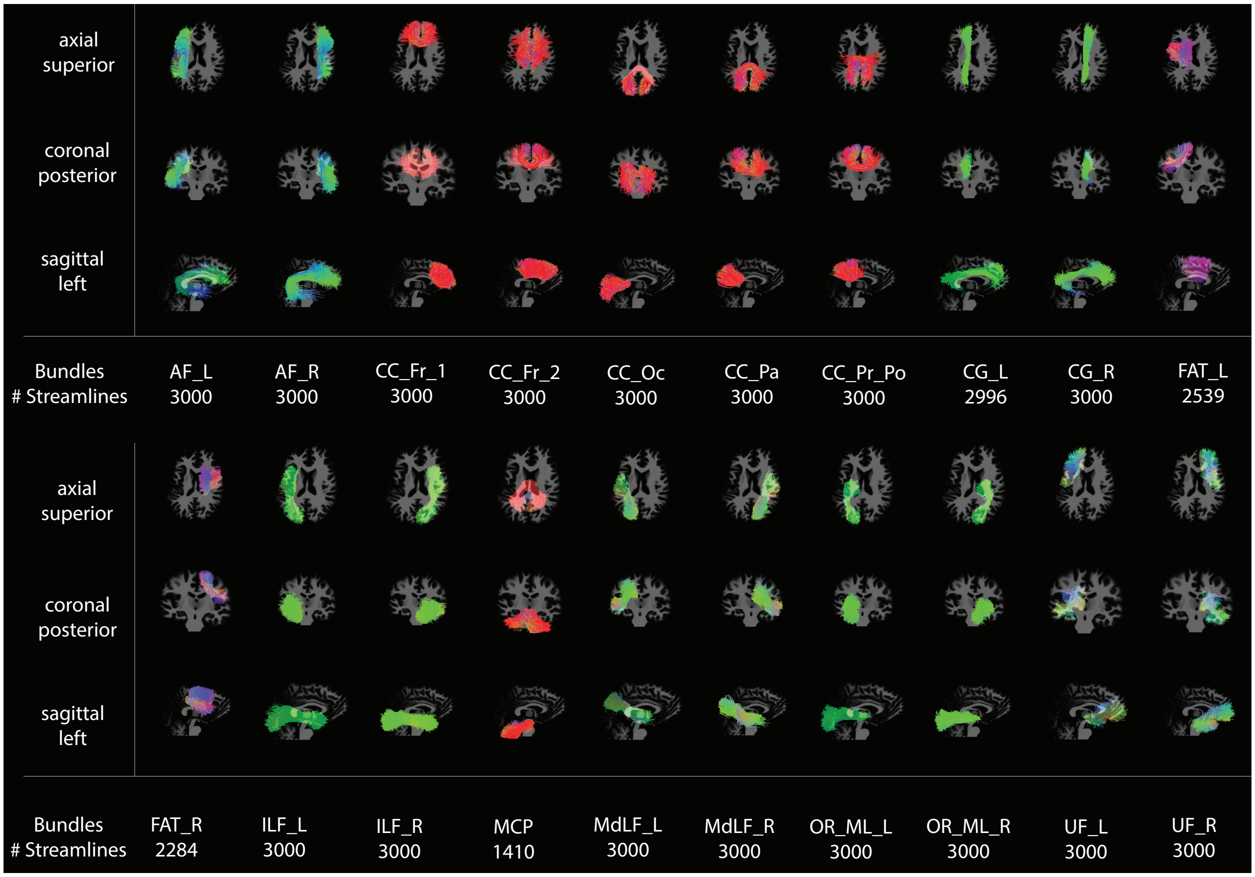


**Fig. S4 | Bundle comparison.** Bundles generated with ATM-Population for a single representative subject.

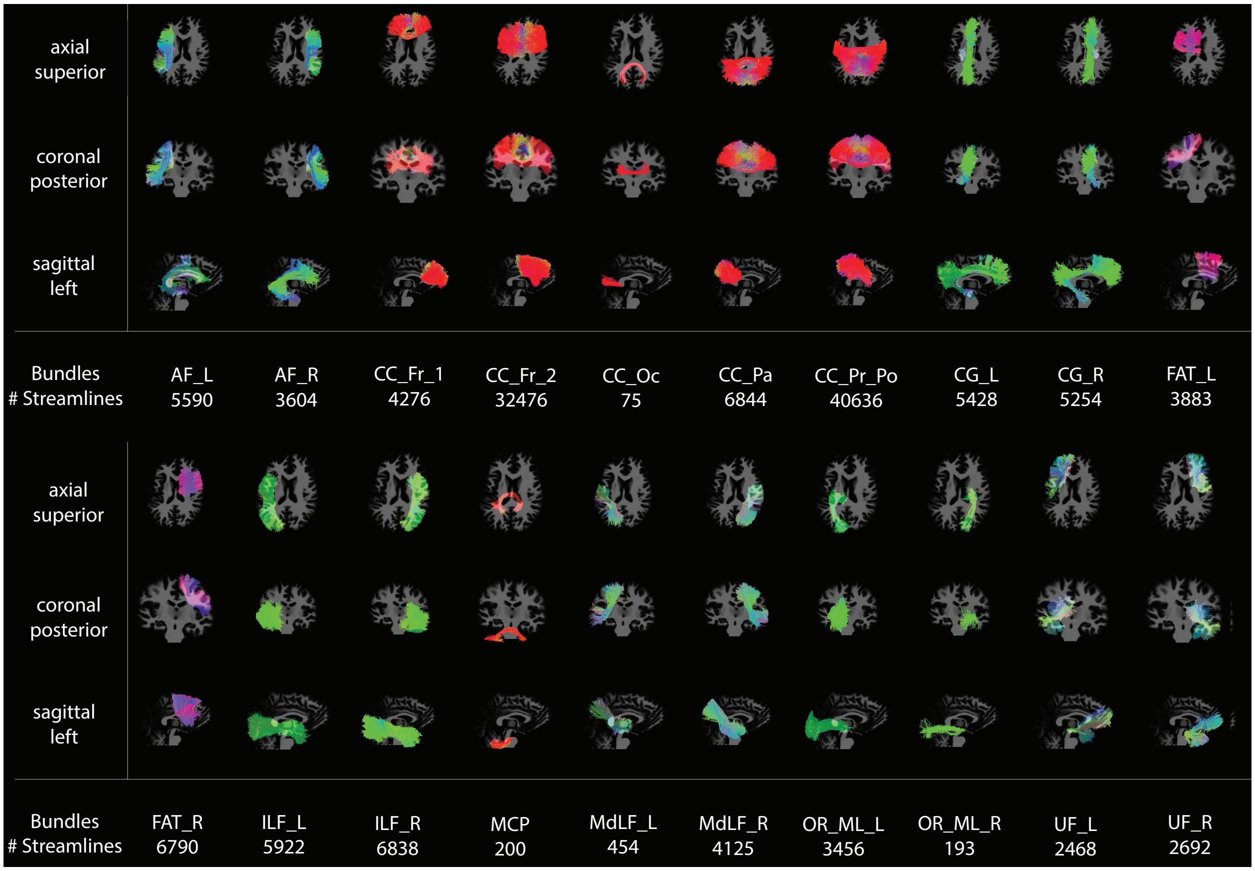


**Fig. S5** | **Bundle comparison.** Bundles generated with MRtrix and BundleSeg for a single representative subject.

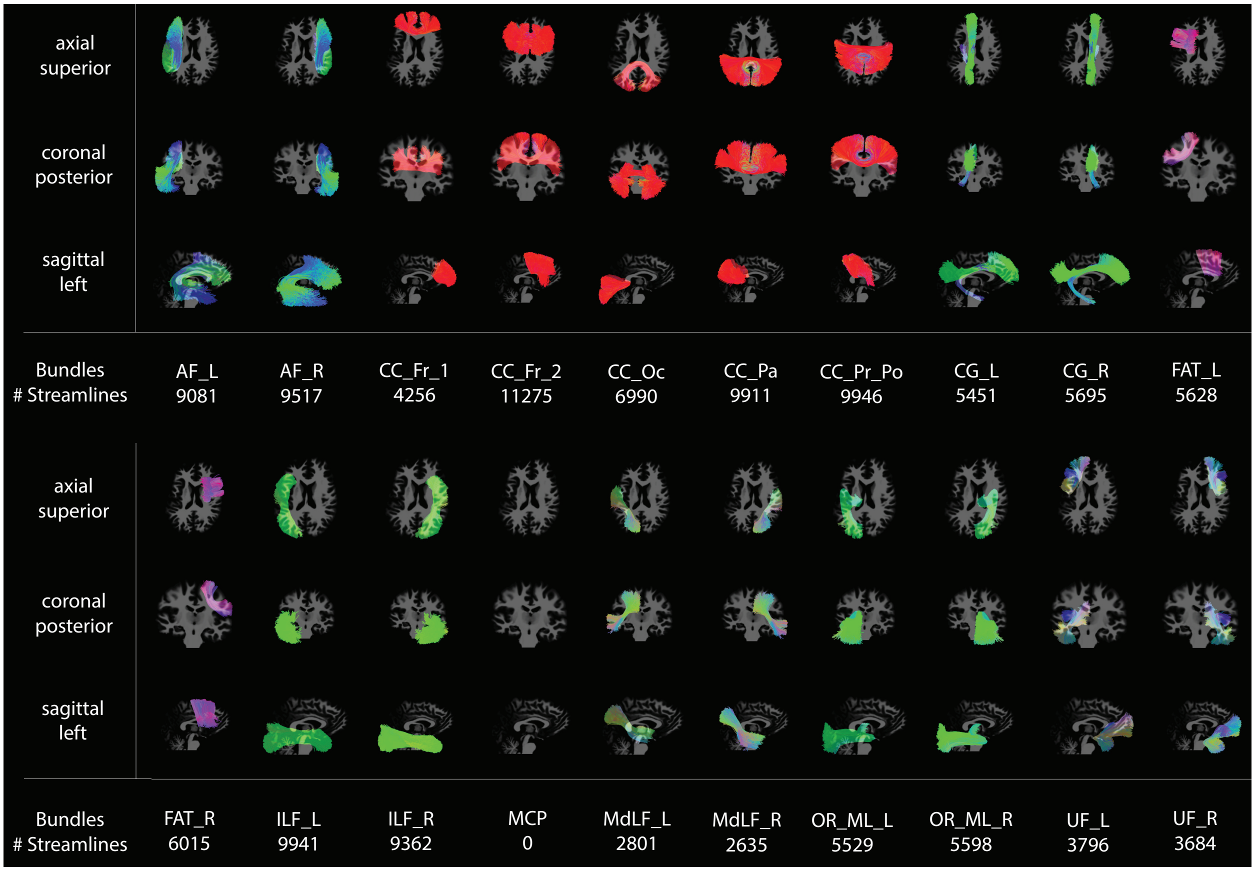


**Fig. S6** | **Bundle comparison.** Bundles generated with SCIL WM atlas warping for a single representative subject.

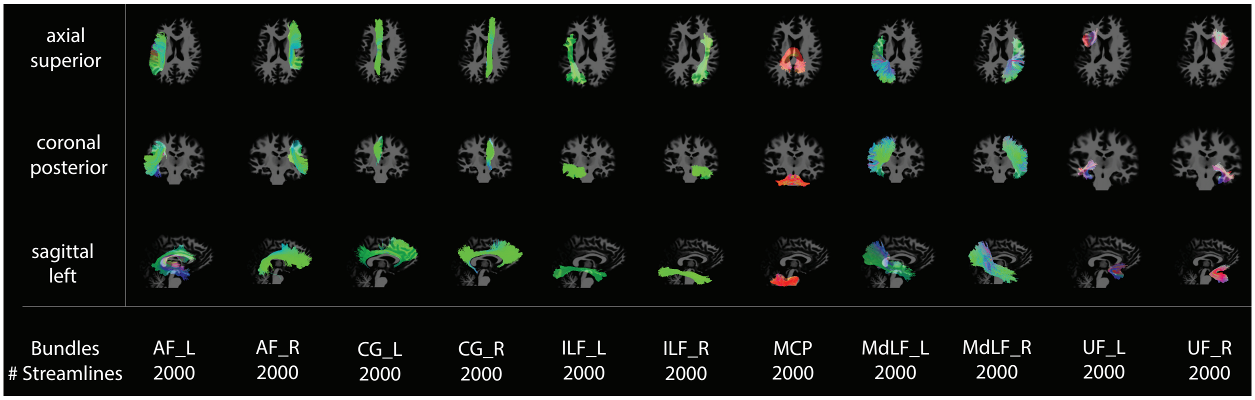


**Fig. S7 | Bundle comparison.** Bundles generated with TractSeg for a single representative subject.

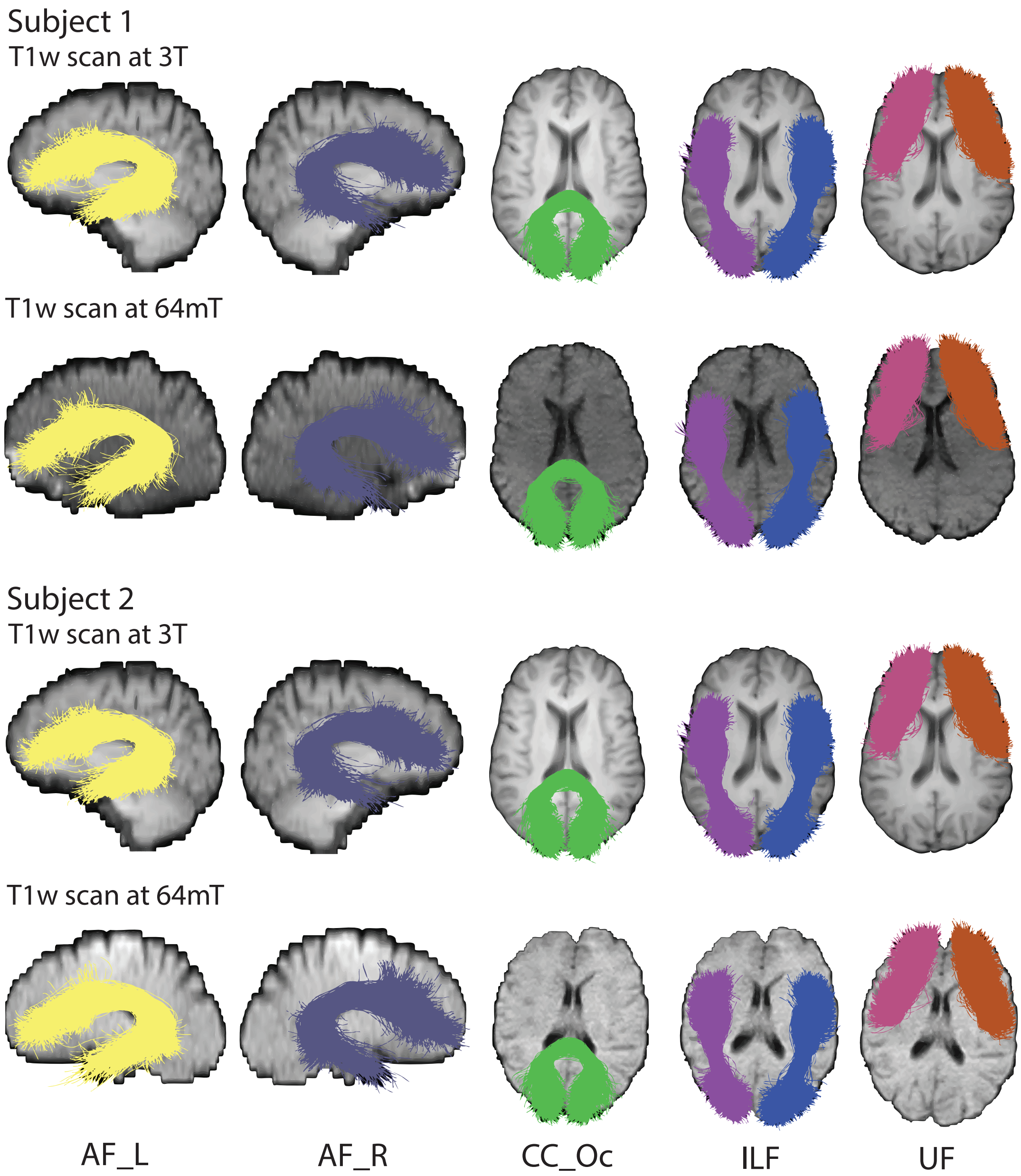


**Fig. S8** | **Clinical scans.** Bundles generated with ATM for clincal T1w scans acquired at 3T with 1.5 mm × 1.5 mm × 5 mm resolution and at 64 mT with 1.6 mm × 1.6 mm × 5 mm resolution.

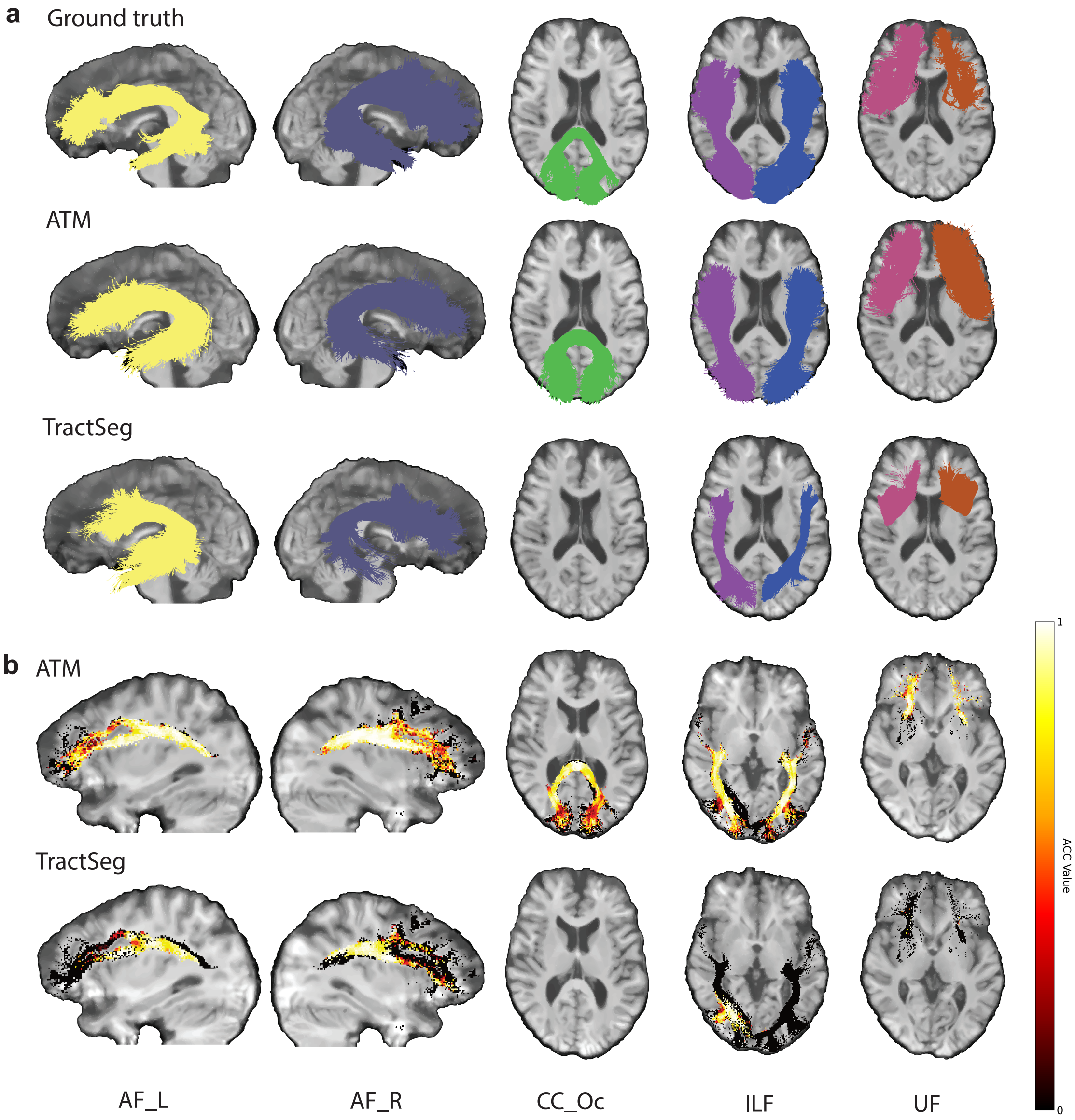


**Fig. S9 | Bundle comparison and orientation alignment.** **a**, Qualitative comparison of representative bundles reconstructed with ATM and TractSeg. **b**, Angular correlation coefficients (ACCs) computed between the tract orientation density maps of the ground truth and generated streamlines. TractSeg does not generate the CC_Oc bundle, as this bundle was not part of its training set. (TractoInferno Subject ID: 1135)

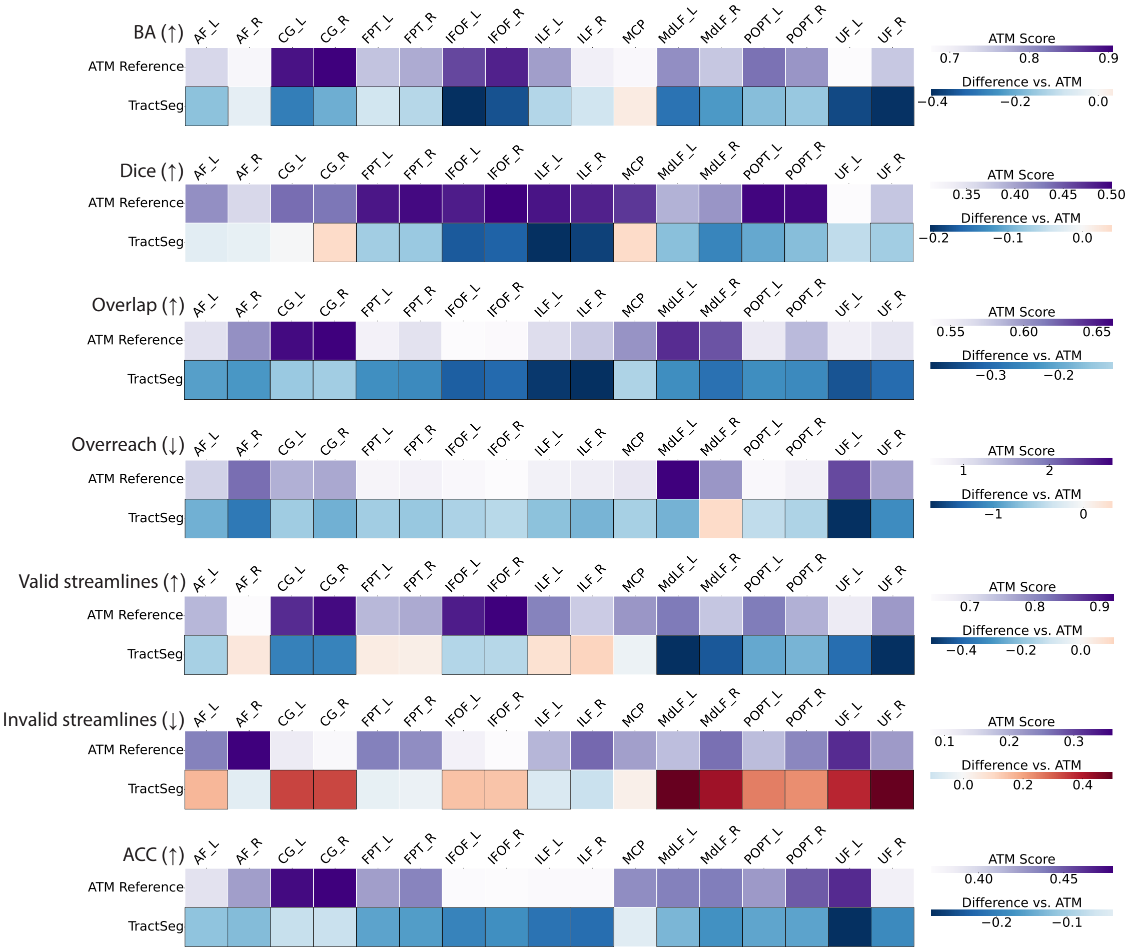


**Fig. S10 | Similarity and coverage of bundles generated with ATM and TractSeg.** The first row of each heatmap shows the ATM values as the baseline. The subsequent row displays the paired performance differences of TractSeg relative to this baseline. For each metric, the arrow indicates preference: ↑ higher is better and ↓ lower is better. Cells outlined with squares indicate statistically significant differences.

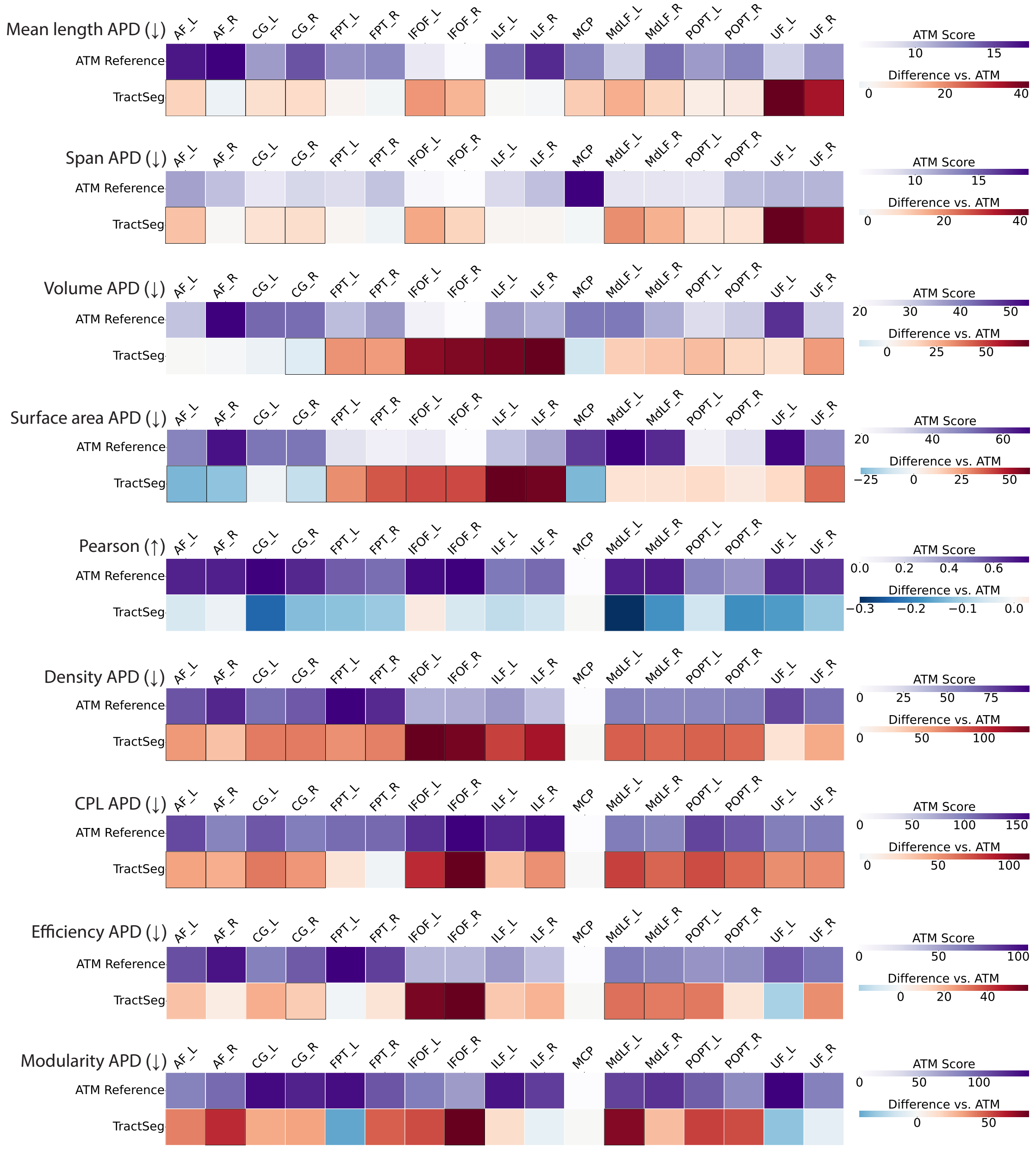


**Fig. S11 | Geometry and connectomics of bundles generated with ATM and TractSeg.** The first row of each heatmap shows the ATM values as the baseline. The subsequent row displays the paired performance differences of TractSeg relative to this baseline. For each metric, the arrow indicates preference: ↑ higher is better and ↓ lower is better. Cells outlined in squares indicate statistically significant differences.

**Tab. S2 | Similarity, coverage, and validity of individual bundles generated with ATM.** Values are presented as mean ± standard deviation. For each metric, the arrow indicates preference: ↑ higher is better, ↓ lower is better.

| Bundles | BA (↑) | Coverage | | | Validity | | ACC (↑) |
|---|---|---|---|---|---|---|---|
| | | Dice (↑) | Overlap (↑) | Overreach (↓) | VS (↑) | IS (↓) | |
| AF_L | 0.7365 ± 0.1000 | 0.4149 ± 0.0540 | 0.5621 ± 0.0817 | 1.2206 ± 0.6170 | 0.7524 ± 0.1310 | 0.2476 ± 0.1310 | 0.3934 ± 0.0665 |
| AF_R | 0.6876 ± 0.1392 | 0.3607 ± 0.0878 | 0.6036 ± 0.0939 | 2.0266 ± 1.2352 | 0.6403 ± 0.2039 | 0.3597 ± 0.2039 | 0.4239 ± 0.0667 |
| CC_Fr_1 | 0.8637 ± 0.1135 | 0.5175 ± 0.0585 | 0.5203 ± 0.0983 | 0.4976 ± 0.2881 | 0.8855 ± 0.1199 | 0.1145 ± 0.1199 | 0.3742 ± 0.0819 |
| CC_Fr_2 | 0.8498 ± 0.0653 | 0.4857 ± 0.0509 | 0.4369 ± 0.0965 | 0.3497 ± 0.2158 | 0.9158 ± 0.0717 | 0.0842 ± 0.0717 | 0.3418 ± 0.0737 |
| CC_Oc | 0.8516 ± 0.0663 | 0.5287 ± 0.0489 | 0.5758 ± 0.0635 | 0.6150 ± 0.2440 | 0.8278 ± 0.0885 | 0.1722 ± 0.0885 | 0.3932 ± 0.0442 |
| CC_Pa | 0.9420 ± 0.0178 | 0.5848 ± 0.0281 | 0.5633 ± 0.0830 | 0.3592 ± 0.1676 | 0.9567 ± 0.0216 | 0.0433 ± 0.0216 | 0.4269 ± 0.0658 |
| CC_Pr_Po | 0.9355 ± 0.0341 | 0.5560 ± 0.0428 | 0.4958 ± 0.0727 | 0.2805 ± 0.0942 | 0.9615 ± 0.0162 | 0.0385 ± 0.0162 | 0.3774 ± 0.0560 |
| CG_L | 0.8914 ± 0.0910 | 0.4382 ± 0.0831 | 0.6571 ± 0.0639 | 1.5003 ± 0.8744 | 0.8824 ± 0.1431 | 0.1176 ± 0.1431 | 0.4751 ± 0.0543 |
| CG_R | 0.9055 ± 0.0552 | 0.4331 ± 0.0765 | 0.6610 ± 0.0592 | 1.5706 ± 1.0833 | 0.9123 ± 0.0717 | 0.0877 ± 0.0717 | 0.4788 ± 0.0645 |
| FAT_L | 0.8702 ± 0.0627 | 0.5504 ± 0.0448 | 0.5753 ± 0.0711 | 0.5303 ± 0.2840 | 0.8826 ± 0.0815 | 0.1174 ± 0.0815 | 0.4272 ± 0.0533 |
| FAT_R | 0.8480 ± 0.0516 | 0.5239 ± 0.0432 | 0.5634 ± 0.0750 | 0.5941 ± 0.2330 | 0.8675 ± 0.0587 | 0.1325 ± 0.0587 | 0.4184 ± 0.0599 |
| FPT_L | 0.7563 ± 0.1323 | 0.4870 ± 0.0659 | 0.5472 ± 0.0730 | 0.7568 ± 0.4906 | 0.7511 ± 0.1355 | 0.2489 ± 0.1355 | 0.4240 ± 0.0639 |
| FPT_R | 0.7769 ± 0.1017 | 0.4946 ± 0.0761 | 0.5611 ± 0.0575 | 0.8062 ± 0.6539 | 0.7650 ± 0.1150 | 0.2350 ± 0.1150 | 0.4354 ± 0.0442 |
| IFOF_L | 0.8548 ± 0.0552 | 0.4832 ± 0.0316 | 0.5364 ± 0.0530 | 0.6971 ± 0.2514 | 0.8963 ± 0.0486 | 0.1037 ± 0.0486 | 0.3730 ± 0.0422 |
| IFOF_R | 0.8811 ± 0.0411 | 0.5004 ± 0.0236 | 0.5386 ± 0.0454 | 0.6181 ± 0.1696 | 0.9217 ± 0.0406 | 0.0783 ± 0.0406 | 0.3720 ± 0.0347 |
| ILF_L | 0.7876 ± 0.1054 | 0.4885 ± 0.0686 | 0.5646 ± 0.0707 | 0.8207 ± 0.5547 | 0.8086 ± 0.1227 | 0.1914 ± 0.1227 | 0.3742 ± 0.0519 |
| ILF_R | 0.7006 ± 0.2435 | 0.4799 ± 0.0684 | 0.5771 ± 0.0882 | 0.8951 ± 0.5654 | 0.7285 ± 0.2502 | 0.2715 ± 0.2502 | 0.3748 ± 0.0689 |
| MCP | 0.6850 ± 0.1771 | 0.4674 ± 0.0662 | 0.6025 ± 0.1354 | 0.9755 ± 0.3588 | 0.7861 ± 0.1311 | 0.2139 ± 0.1311 | 0.4316 ± 0.1100 |
| MdLF_L | 0.8025 ± 0.1923 | 0.3904 ± 0.1373 | 0.6433 ± 0.1247 | 2.7284 ± 4.9993 | 0.8178 ± 0.2170 | 0.1822 ± 0.2170 | 0.4371 ± 0.0945 |
| MdLF_R | 0.7530 ± 0.2253 | 0.4104 ± 0.1407 | 0.6292 ± 0.1647 | 1.7125 ± 1.1728 | 0.7355 ± 0.2440 | 0.2645 ± 0.2440 | 0.4381 ± 0.1223 |
| OR_ML_L | 0.9234 ± 0.0521 | 0.4670 ± 0.0689 | 0.6785 ± 0.0812 | 1.3203 ± 0.6718 | 0.9218 ± 0.0610 | 0.0782 ± 0.0610 | 0.4705 ± 0.0671 |
| OR_ML_R | 0.8520 ± 0.1234 | 0.3730 ± 0.1016 | 0.6928 ± 0.1032 | 2.2754 ± 1.1133 | 0.8479 ± 0.1626 | 0.1521 ± 0.1626 | 0.4754 ± 0.0725 |
| POPT_L | 0.8267 ± 0.0906 | 0.4973 ± 0.0628 | 0.5550 ± 0.0760 | 0.6965 ± 0.2918 | 0.8161 ± 0.0938 | 0.1839 ± 0.0938 | 0.4256 ± 0.0652 |
| POPT_R | 0.7955 ± 0.0784 | 0.4955 ± 0.0762 | 0.5842 ± 0.0783 | 0.8341 ± 0.5008 | 0.7581 ± 0.1239 | 0.2419 ± 0.1239 | 0.4496 ± 0.0664 |
| PYT_L | 0.8664 ± 0.0944 | 0.5673 ± 0.0345 | 0.5306 ± 0.0577 | 0.3379 ± 0.1038 | 0.8817 ± 0.0573 | 0.1183 ± 0.0573 | 0.4097 ± 0.0488 |
| PYT_R | 0.8661 ± 0.0857 | 0.5698 ± 0.0342 | 0.5360 ± 0.0446 | 0.3459 ± 0.0931 | 0.8913 ± 0.0514 | 0.1087 ± 0.0514 | 0.4169 ± 0.0405 |
| SLF_L | 0.5883 ± 0.1786 | 0.3268 ± 0.1011 | 0.5707 ± 0.1311 | 2.1464 ± 1.0886 | 0.5322 ± 0.1935 | 0.4678 ± 0.1935 | 0.3934 ± 0.0924 |
| SLF_R | 0.6989 ± 0.1487 | 0.4024 ± 0.1100 | 0.6033 ± 0.0863 | 1.6959 ± 1.1546 | 0.6568 ± 0.2123 | 0.3432 ± 0.2123 | 0.4090 ± 0.0634 |
| UF_L | 0.6763 ± 0.2320 | 0.3121 ± 0.1300 | 0.5507 ± 0.1630 | 2.2471 ± 1.1113 | 0.6790 ± 0.2396 | 0.3210 ± 0.2396 | 0.4644 ± 0.0975 |
| UF_R | 0.7524 ± 0.1792 | 0.3761 ± 0.1096 | 0.5588 ± 0.1258 | 1.5944 ± 0.9047 | 0.7814 ± 0.1933 | 0.2186 ± 0.1933 | 0.3827 ± 0.1084 |

**Tab. S3 | Similarity, coverage, and validity of population bundles generated with ATM.** Values are presented as mean ± standard deviation. For each metric, the arrow indicates preference: ↑ higher is better, ↓ lower is better.

| Bundles | BA (↑) | Coverage | | | Validity | | ACC (↑) |
|---|---|---|---|---|---|---|---|
| | | Dice (↑) | Overlap (↑) | Overreach (↓) | VS (↑) | IS (↓) | |
| AF_L | 0.4972 ± 0.2037 | 0.3491 ± 0.0681 | 0.4503 ± 0.0791 | 1.2223 ± 0.6353 | 0.5695 ± 0.2216 | 0.4305 ± 0.2216 | 0.2573 ± 0.0555 |
| AF_R | 0.5316 ± 0.1828 | 0.3058 ± 0.0862 | 0.5436 ± 0.1026 | 2.4976 ± 1.7982 | 0.5490 ± 0.2407 | 0.4510 ± 0.2407 | 0.3118 ± 0.0679 |
| CC_Fr_1 | 0.7402 ± 0.2111 | 0.4339 ± 0.1301 | 0.4257 ± 0.1648 | 0.4769 ± 0.3904 | 0.7461 ± 0.2570 | 0.2539 ± 0.2570 | 0.2626 ± 0.1043 |
| CC_Fr_2 | 0.7878 ± 0.1797 | 0.4118 ± 0.1323 | 0.3708 ± 0.1521 | 0.3579 ± 0.2682 | 0.8793 ± 0.1360 | 0.1207 ± 0.1360 | 0.2387 ± 0.0986 |
| CC_Oc | 0.5739 ± 0.2737 | 0.4081 ± 0.0733 | 0.3907 ± 0.0903 | 0.5208 ± 0.2446 | 0.7162 ± 0.2214 | 0.2838 ± 0.2214 | 0.2116 ± 0.0553 |
| CC_Pa | 0.8306 ± 0.0778 | 0.4741 ± 0.0512 | 0.4119 ± 0.0744 | 0.3178 ± 0.1560 | 0.9318 ± 0.0658 | 0.0682 ± 0.0658 | 0.2627 ± 0.0552 |
| CC_Pr_Po | 0.8171 ± 0.1095 | 0.4509 ± 0.0613 | 0.3676 ± 0.0730 | 0.2535 ± 0.1144 | 0.9162 ± 0.0801 | 0.0838 ± 0.0801 | 0.2372 ± 0.0476 |
| CG_L | 0.7453 ± 0.2159 | 0.3804 ± 0.0741 | 0.5076 ± 0.0799 | 1.2862 ± 0.7755 | 0.7597 ± 0.2700 | 0.2403 ± 0.2700 | 0.3070 ± 0.0587 |
| CG_R | 0.7849 ± 0.1907 | 0.3860 ± 0.0793 | 0.5479 ± 0.1191 | 1.3903 ± 0.8518 | 0.8092 ± 0.1876 | 0.1908 ± 0.1876 | 0.3343 ± 0.0999 |
| FAT_L | 0.6981 ± 0.1695 | 0.4439 ± 0.1202 | 0.4178 ± 0.1325 | 0.4142 ± 0.2182 | 0.8013 ± 0.1198 | 0.1987 ± 0.1198 | 0.2543 ± 0.0833 |
| FAT_R | 0.7079 ± 0.1700 | 0.4289 ± 0.1155 | 0.4306 ± 0.1525 | 0.5111 ± 0.2746 | 0.8148 ± 0.1224 | 0.1852 ± 0.1224 | 0.2626 ± 0.1002 |
| FPT_L | 0.5383 ± 0.1815 | 0.4203 ± 0.0658 | 0.4774 ± 0.0762 | 0.8403 ± 0.4379 | 0.6112 ± 0.1760 | 0.3888 ± 0.1760 | 0.3066 ± 0.0595 |
| FPT_R | 0.5186 ± 0.1978 | 0.4226 ± 0.0708 | 0.4834 ± 0.0625 | 0.8942 ± 0.6174 | 0.5802 ± 0.2092 | 0.4198 ± 0.2092 | 0.3120 ± 0.0520 |
| IFOF_L | 0.5334 ± 0.2346 | 0.3883 ± 0.0431 | 0.3914 ± 0.0560 | 0.6302 ± 0.2024 | 0.7586 ± 0.1817 | 0.2414 ± 0.1817 | 0.2276 ± 0.0453 |
| IFOF_R | 0.6637 ± 0.1795 | 0.4203 ± 0.0222 | 0.4422 ± 0.0447 | 0.6623 ± 0.1456 | 0.8098 ± 0.1325 | 0.1902 ± 0.1325 | 0.2593 ± 0.0278 |
| ILF_L | 0.4647 ± 0.2355 | 0.3793 ± 0.0671 | 0.3917 ± 0.0619 | 0.7321 ± 0.4640 | 0.6535 ± 0.2471 | 0.3465 ± 0.2471 | 0.2099 ± 0.0479 |
| ILF_R | 0.5136 ± 0.1872 | 0.3859 ± 0.0478 | 0.4051 ± 0.0534 | 0.7195 ± 0.3312 | 0.6863 ± 0.1998 | 0.3137 ± 0.1998 | 0.2212 ± 0.0353 |
| MCP | 0.2257 ± 0.2003 | 0.2593 ± 0.0900 | 0.3470 ± 0.1217 | 1.3768 ± 0.5541 | 0.2722 ± 0.2075 | 0.7278 ± 0.2075 | 0.1946 ± 0.0895 |
| MdLF_L | 0.5507 ± 0.2734 | 0.3272 ± 0.1194 | 0.4919 ± 0.1000 | 2.3893 ± 2.9293 | 0.6272 ± 0.3094 | 0.3728 ± 0.3094 | 0.2748 ± 0.0699 |
| MdLF_R | 0.5814 ± 0.2460 | 0.3358 ± 0.1260 | 0.5265 ± 0.1133 | 3.1412 ± 5.0060 | 0.6132 ± 0.2908 | 0.3868 ± 0.2908 | 0.3050 ± 0.0732 |
| OR_ML_L | 0.7133 ± 0.1918 | 0.3732 ± 0.0820 | 0.4711 ± 0.1067 | 1.1130 ± 0.5520 | 0.8134 ± 0.1872 | 0.1866 ± 0.1872 | 0.2679 ± 0.0742 |
| OR_ML_R | 0.6056 ± 0.2224 | 0.2723 ± 0.0804 | 0.4588 ± 0.1184 | 2.4736 ± 2.4420 | 0.6605 ± 0.2243 | 0.3395 ± 0.2243 | 0.2674 ± 0.0876 |
| POPT_L | 0.5037 ± 0.1883 | 0.3901 ± 0.1069 | 0.4100 ± 0.1145 | 0.6741 ± 0.3649 | 0.6473 ± 0.2115 | 0.3527 ± 0.2115 | 0.2683 ± 0.0804 |
| POPT_R | 0.4539 ± 0.2014 | 0.3854 ± 0.1132 | 0.4226 ± 0.1190 | 0.7927 ± 0.5398 | 0.5876 ± 0.2328 | 0.4124 ± 0.2328 | 0.2756 ± 0.0815 |
| PYT_L | 0.5486 ± 0.1989 | 0.4521 ± 0.1007 | 0.4086 ± 0.1027 | 0.3651 ± 0.1428 | 0.7075 ± 0.2088 | 0.2925 ± 0.2088 | 0.2664 ± 0.0701 |
| PYT_R | 0.6477 ± 0.2030 | 0.4927 ± 0.0491 | 0.4653 ± 0.0682 | 0.4194 ± 0.1368 | 0.8091 ± 0.1299 | 0.1909 ± 0.1299 | 0.3091 ± 0.0455 |
| SLF_L | 0.4941 ± 0.2052 | 0.2696 ± 0.1062 | 0.5140 ± 0.1481 | 3.0548 ± 3.0817 | 0.5111 ± 0.2381 | 0.4889 ± 0.2381 | 0.2891 ± 0.0856 |
| SLF_R | 0.5760 ± 0.2273 | 0.3378 ± 0.1159 | 0.4722 ± 0.1280 | 1.7756 ± 1.7986 | 0.6202 ± 0.2685 | 0.3798 ± 0.2685 | 0.2680 ± 0.0705 |
| UF_L | 0.3520 ± 0.1894 | 0.2374 ± 0.1007 | 0.4278 ± 0.0899 | 3.0130 ± 2.2913 | 0.4130 ± 0.2245 | 0.5870 ± 0.2245 | 0.2277 ± 0.0527 |
| UF_R | 0.4706 ± 0.1851 | 0.2848 ± 0.0984 | 0.4681 ± 0.0800 | 2.6749 ± 3.0466 | 0.5408 ± 0.1865 | 0.4592 ± 0.1865 | 0.2595 ± 0.0524 |

**Tab. S4 | Similarity, coverage, and validity of bundles generated from MRtrix with BundleSeg.** Values are presented as mean ± standard deviation. For each metric, the arrow indicates preference: ↑ higher is better, ↓ lower is better.

| Bundles | BA (↑) | Coverage | | | Validity | | ACC (↑) |
|---|---|---|---|---|---|---|---|
| | | Dice (↑) | Overlap (↑) | Overreach (↓) | VS (↑) | IS (↓) | |
| AF_L | 0.3683 ± 0.1650 | 0.4492 ± 0.0846 | 0.7046 ± 0.1479 | 1.6085 ± 1.1178 | 0.5412 ± 0.2079 | 0.4588 ± 0.2079 | 0.5826 ± 0.1174 |
| AF_R | 0.3163 ± 0.1808 | 0.3656 ± 0.1204 | 0.7238 ± 0.1652 | 2.6595 ± 1.8034 | 0.4552 ± 0.1903 | 0.5448 ± 0.1903 | 0.5996 ± 0.1350 |
| CC_Fr_1 | 0.8905 ± 0.0640 | 0.5636 ± 0.1282 | 0.7193 ± 0.1915 | 0.8672 ± 0.7374 | 0.9125 ± 0.0768 | 0.0875 ± 0.0768 | 0.6034 ± 0.1558 |
| CC_Fr_2 | 0.8175 ± 0.1572 | 0.5399 ± 0.1343 | 0.7068 ± 0.1587 | 1.0329 ± 0.7882 | 0.6894 ± 0.3198 | 0.3106 ± 0.3198 | 0.6098 ± 0.1351 |
| CC_Oc | 0.3968 ± 0.1495 | 0.3944 ± 0.1252 | 0.2930 ± 0.1160 | 0.1356 ± 0.1137 | 0.7922 ± 0.1725 | 0.2078 ± 0.1725 | 0.2634 ± 0.1008 |
| CC_Pa | 0.7782 ± 0.0762 | 0.6095 ± 0.0585 | 0.7110 ± 0.1243 | 0.6534 ± 0.4556 | 0.8724 ± 0.0833 | 0.1276 ± 0.0833 | 0.6111 ± 0.0982 |
| CC_Pr_Po | 0.7370 ± 0.0973 | 0.5371 ± 0.0734 | 0.7707 ± 0.1131 | 1.1416 ± 0.4426 | 0.7432 ± 0.3368 | 0.2568 ± 0.3368 | 0.6660 ± 0.1010 |
| CG_L | 0.6219 ± 0.1240 | 0.3017 ± 0.0834 | 0.8114 ± 0.1543 | 4.1982 ± 2.3738 | 0.5804 ± 0.1379 | 0.4196 ± 0.1379 | 0.6573 ± 0.1217 |
| CG_R | 0.6737 ± 0.1189 | 0.2928 ± 0.0747 | 0.8484 ± 0.1875 | 4.7850 ± 3.5378 | 0.5914 ± 0.1594 | 0.4086 ± 0.1594 | 0.6842 ± 0.1489 |
| FAT_L | 0.8459 ± 0.0933 | 0.5357 ± 0.1352 | 0.7262 ± 0.1840 | 0.9707 ± 0.5377 | 0.8286 ± 0.1536 | 0.1714 ± 0.1536 | 0.6109 ± 0.1535 |
| FAT_R | 0.8614 ± 0.0703 | 0.5378 ± 0.1085 | 0.7672 ± 0.1668 | 1.1163 ± 0.5926 | 0.8397 ± 0.1240 | 0.1603 ± 0.1240 | 0.6436 ± 0.1404 |
| FPT_L | 0.5240 ± 0.1923 | 0.5301 ± 0.1020 | 0.5573 ± 0.1345 | 0.5892 ± 0.4804 | 0.7561 ± 0.1977 | 0.2439 ± 0.1977 | 0.4853 ± 0.1165 |
| FPT_R | 0.5802 ± 0.1641 | 0.5528 ± 0.1000 | 0.6328 ± 0.1372 | 0.7238 ± 0.6504 | 0.8074 ± 0.1424 | 0.1926 ± 0.1424 | 0.5499 ± 0.1155 |
| IFOF_L | 0.3206 ± 0.1565 | 0.4141 ± 0.0414 | 0.3397 ± 0.0817 | 0.2832 ± 0.1592 | 0.7336 ± 0.2674 | 0.2664 ± 0.2674 | 0.2949 ± 0.0636 |
| IFOF_R | 0.3830 ± 0.1789 | 0.4403 ± 0.0476 | 0.3791 ± 0.0815 | 0.3304 ± 0.1778 | 0.7650 ± 0.2371 | 0.2350 ± 0.2371 | 0.3260 ± 0.0657 |
| ILF_L | 0.6713 ± 0.1454 | 0.5047 ± 0.1002 | 0.8328 ± 0.1312 | 1.6741 ± 1.1211 | 0.7781 ± 0.1425 | 0.2219 ± 0.1425 | 0.6674 ± 0.0964 |
| ILF_R | 0.6777 ± 0.1254 | 0.4990 ± 0.0945 | 0.8614 ± 0.1290 | 1.7669 ± 0.9854 | 0.7398 ± 0.1306 | 0.2602 ± 0.1306 | 0.6902 ± 0.0943 |
| MCP | 0.5711 ± 0.3029 | 0.5121 ± 0.1283 | 0.5946 ± 0.1861 | 0.6924 ± 0.4555 | 0.8071 ± 0.1948 | 0.1929 ± 0.1948 | 0.5006 ± 0.1588 |
| MdLF_L | 0.6611 ± 0.2661 | 0.4130 ± 0.1593 | 0.7546 ± 0.1549 | 3.0138 ± 3.8191 | 0.6827 ± 0.2972 | 0.3173 ± 0.2972 | 0.6176 ± 0.1275 |
| MdLF_R | 0.7477 ± 0.1913 | 0.4072 ± 0.1748 | 0.8202 ± 0.1388 | 4.9458 ± 7.3139 | 0.7009 ± 0.2969 | 0.2991 ± 0.2969 | 0.6708 ± 0.1112 |
| OR_ML_L | 0.9042 ± 0.0721 | 0.5308 ± 0.0847 | 0.7879 ± 0.0908 | 1.2713 ± 0.6414 | 0.9267 ± 0.0670 | 0.0733 ± 0.0670 | 0.6592 ± 0.0698 |
| OR_ML_R | 0.8524 ± 0.1013 | 0.4447 ± 0.1344 | 0.8207 ± 0.1111 | 2.6992 ± 3.0106 | 0.8444 ± 0.1495 | 0.1556 ± 0.1495 | 0.6936 ± 0.0909 |
| POPT_L | 0.5587 ± 0.1489 | 0.5202 ± 0.0501 | 0.5473 ± 0.1021 | 0.5738 ± 0.3610 | 0.7563 ± 0.1735 | 0.2437 ± 0.1735 | 0.4826 ± 0.0874 |
| POPT_R | 0.5262 ± 0.1475 | 0.5127 ± 0.0628 | 0.5632 ± 0.1006 | 0.7098 ± 0.6573 | 0.6628 ± 0.2163 | 0.3372 ± 0.2163 | 0.4978 ± 0.0863 |
| PYT_L | 0.7241 ± 0.1753 | 0.6260 ± 0.0478 | 0.6521 ± 0.1193 | 0.4202 ± 0.1951 | 0.9008 ± 0.1160 | 0.0992 ± 0.1160 | 0.5729 ± 0.1020 |
| PYT_R | 0.6987 ± 0.1566 | 0.6345 ± 0.0359 | 0.6645 ± 0.1067 | 0.4277 ± 0.2162 | 0.8891 ± 0.0832 | 0.1109 ± 0.0832 | 0.5845 ± 0.0912 |
| SLF_L | 0.4285 ± 0.2163 | 0.3149 ± 0.1284 | 0.7706 ± 0.2017 | 4.2206 ± 4.6380 | 0.4369 ± 0.2183 | 0.5631 ± 0.2183 | 0.6396 ± 0.1649 |
| SLF_R | 0.5438 ± 0.2388 | 0.3932 ± 0.1662 | 0.7881 ± 0.2052 | 3.0247 ± 3.0350 | 0.5999 ± 0.2983 | 0.4001 ± 0.2983 | 0.6472 ± 0.1661 |
| UF_L | 0.6769 ± 0.2283 | 0.3499 ± 0.1699 | 0.7080 ± 0.2067 | 3.5978 ± 3.6483 | 0.6795 ± 0.2525 | 0.3205 ± 0.2525 | 0.5641 ± 0.1712 |
| UF_R | 0.7499 ± 0.1695 | 0.3942 ± 0.1262 | 0.7307 ± 0.1888 | 2.6260 ± 2.5430 | 0.7690 ± 0.1871 | 0.2310 ± 0.1871 | 0.5765 ± 0.1529 |

**Tab. S5 | Similarity, coverage, and validity of population bundles generated with SCIL WM atlas warping.** Values are presented as mean ± standard deviation. For each metric, the arrow indicates preference: ↑ higher is better, ↓ lower is better.

| Bundles | BA (↑) | Coverage | | | Validity | | ACC (↑) |
|---|---|---|---|---|---|---|---|
| | | Dice (↑) | Overlap (↑) | Overreach (↓) | VS (↑) | IS (↓) | |
| AF_L | 0.6292 ± 0.1495 | 0.3523 ± 0.0667 | 0.5329 ± 0.0711 | 1.6392 ± 0.8364 | 0.5954 ± 0.2042 | 0.4046 ± 0.2042 | 0.3077 ± 0.0644 |
| AF_R | 0.5541 ± 0.1369 | 0.2776 ± 0.0924 | 0.5788 ± 0.0928 | 3.2946 ± 2.3145 | 0.4686 ± 0.1955 | 0.5314 ± 0.1955 | 0.3224 ± 0.0892 |
| CC_Fr_1 | 0.2417 ± 0.2213 | 0.2709 ± 0.0329 | 0.2052 ± 0.0450 | 0.2987 ± 0.1531 | 0.4619 ± 0.2753 | 0.5381 ± 0.2753 | 0.1032 ± 0.0406 |
| CC_Fr_2 | 0.5255 ± 0.1936 | 0.3143 ± 0.0540 | 0.2615 ± 0.0509 | 0.4174 ± 0.2474 | 0.7674 ± 0.1836 | 0.2326 ± 0.1836 | 0.1560 ± 0.0423 |
| CC_Oc | 0.5984 ± 0.2261 | 0.3213 ± 0.0502 | 0.2582 ± 0.0470 | 0.3502 ± 0.1354 | 0.7690 ± 0.1932 | 0.2310 ± 0.1932 | 0.1587 ± 0.0354 |
| CC_Pa | 0.7669 ± 0.1166 | 0.4004 ± 0.0415 | 0.3813 ± 0.0481 | 0.5327 ± 0.2108 | 0.8209 ± 0.0726 | 0.1791 ± 0.0726 | 0.2516 ± 0.0398 |
| CC_Pr_Po | 0.7306 ± 0.1673 | 0.3582 ± 0.0624 | 0.3062 ± 0.0510 | 0.4139 ± 0.1517 | 0.7544 ± 0.1748 | 0.2456 ± 0.1748 | 0.2127 ± 0.0451 |
| CG_L | 0.4952 ± 0.2307 | 0.2987 ± 0.0799 | 0.3872 ± 0.0640 | 1.3788 ± 0.7905 | 0.5585 ± 0.2572 | 0.4415 ± 0.2572 | 0.2324 ± 0.0596 |
| CG_R | 0.5821 ± 0.1856 | 0.2992 ± 0.0716 | 0.4061 ± 0.0663 | 1.4719 ± 0.9060 | 0.5927 ± 0.2088 | 0.4073 ± 0.2088 | 0.2392 ± 0.0548 |
| FAT_L | 0.4470 ± 0.1651 | 0.1911 ± 0.0550 | 0.1397 ± 0.0367 | 0.3451 ± 0.1503 | 0.7006 ± 0.2469 | 0.2994 ± 0.2469 | 0.0801 ± 0.0295 |
| FAT_R | 0.5194 ± 0.2347 | 0.2456 ± 0.0666 | 0.1896 ± 0.0483 | 0.3685 ± 0.1486 | 0.7825 ± 0.2316 | 0.2175 ± 0.2316 | 0.1118 ± 0.0328 |
| FPT_L | 0.6110 ± 0.1684 | 0.3147 ± 0.0713 | 0.2647 ± 0.0659 | 0.4382 ± 0.2653 | 0.8380 ± 0.1700 | 0.1620 ± 0.1700 | 0.1766 ± 0.0473 |
| FPT_R | 0.6065 ± 0.2053 | 0.3246 ± 0.0761 | 0.2717 ± 0.0568 | 0.4484 ± 0.3410 | 0.8121 ± 0.2032 | 0.1879 ± 0.2032 | 0.1808 ± 0.0403 |
| IFOF_L | 0.6790 ± 0.1889 | 0.4000 ± 0.0352 | 0.4085 ± 0.0548 | 0.6391 ± 0.2119 | 0.8578 ± 0.0951 | 0.1422 ± 0.0951 | 0.2428 ± 0.0470 |
| IFOF_R | 0.7177 ± 0.1820 | 0.4146 ± 0.0332 | 0.4208 ± 0.0465 | 0.6104 ± 0.1384 | 0.8627 ± 0.1090 | 0.1373 ± 0.1090 | 0.2535 ± 0.0404 |
| ILF_L | 0.7083 ± 0.1922 | 0.4029 ± 0.0689 | 0.3942 ± 0.0632 | 0.6187 ± 0.4298 | 0.8509 ± 0.1859 | 0.1491 ± 0.1859 | 0.2217 ± 0.0472 |
| ILF_R | 0.6627 ± 0.1771 | 0.4006 ± 0.0592 | 0.3969 ± 0.0474 | 0.6194 ± 0.3238 | 0.7953 ± 0.1801 | 0.2047 ± 0.1801 | 0.2203 ± 0.0332 |
| MCP | 0.6261 ± 0.2238 | 0.4197 ± 0.0866 | 0.5127 ± 0.1259 | 0.9374 ± 0.4264 | 0.8406 ± 0.1717 | 0.1594 ± 0.1717 | 0.3127 ± 0.0838 |
| MdLF_L | 0.6514 ± 0.2497 | 0.2961 ± 0.1042 | 0.3268 ± 0.0808 | 1.4348 ± 2.0163 | 0.7699 ± 0.2828 | 0.2301 ± 0.2828 | 0.1952 ± 0.0583 |
| MdLF_R | 0.7184 ± 0.2584 | 0.3245 ± 0.1130 | 0.3534 ± 0.0771 | 1.5316 ± 2.4494 | 0.8059 ± 0.2798 | 0.1941 ± 0.2798 | 0.2218 ± 0.0528 |
| OR_ML_L | 0.5774 ± 0.1592 | 0.3178 ± 0.0565 | 0.3937 ± 0.0613 | 1.1509 ± 0.5152 | 0.6314 ± 0.1724 | 0.3686 ± 0.1724 | 0.2297 ± 0.0472 |
| OR_ML_R | 0.4285 ± 0.1937 | 0.2546 ± 0.0889 | 0.4000 ± 0.1145 | 2.2460 ± 2.1154 | 0.4630 ± 0.1867 | 0.5370 ± 0.1867 | 0.2262 ± 0.0831 |
| POPT_L | 0.5974 ± 0.2028 | 0.2622 ± 0.0529 | 0.2208 ± 0.0473 | 0.4660 ± 0.1570 | 0.8217 ± 0.1960 | 0.1783 ± 0.1960 | 0.1561 ± 0.0376 |
| POPT_R | 0.6190 ± 0.1672 | 0.2785 ± 0.0393 | 0.2513 ± 0.0565 | 0.5452 ± 0.2428 | 0.7678 ± 0.1786 | 0.2322 ± 0.1786 | 0.1777 ± 0.0391 |
| PYT_L | 0.8087 ± 0.1652 | 0.3321 ± 0.0650 | 0.2333 ± 0.0573 | 0.1589 ± 0.0673 | 0.9555 ± 0.0836 | 0.0445 ± 0.0836 | 0.1694 ± 0.0462 |
| PYT_R | 0.8449 ± 0.1374 | 0.3365 ± 0.0643 | 0.2359 ± 0.0557 | 0.1547 ± 0.0550 | 0.9738 ± 0.0504 | 0.0262 ± 0.0504 | 0.1689 ± 0.0427 |
| SLF_L | 0.5144 ± 0.1938 | 0.2685 ± 0.0975 | 0.4601 ± 0.0873 | 2.6943 ± 2.5927 | 0.5086 ± 0.2261 | 0.4914 ± 0.2261 | 0.2471 ± 0.0642 |
| SLF_R | 0.4511 ± 0.2167 | 0.2915 ± 0.0947 | 0.3881 ± 0.0661 | 1.7064 ± 1.4795 | 0.5328 ± 0.2714 | 0.4672 ± 0.2714 | 0.1991 ± 0.0526 |
| UF_L | 0.4147 ± 0.2847 | 0.2466 ± 0.1029 | 0.3379 ± 0.0897 | 2.0120 ± 1.7374 | 0.4868 ± 0.3153 | 0.5132 ± 0.3153 | 0.1992 ± 0.0563 |
| UF_R | 0.5148 ± 0.2697 | 0.2806 ± 0.0986 | 0.3496 ± 0.0690 | 1.6912 ± 2.0572 | 0.5681 ± 0.3401 | 0.4319 ± 0.3401 | 0.2092 ± 0.0535 |

**Tab. S6 | Similarity, coverage, and validity of bundles generated with TractSeg.** Values are presented as mean ± standard deviation. For each metric, the arrow indicates preference: ↑ higher is better, ↓ lower is better.

| Bundles | BA (↑) | | Coverage | | Validity | | ACC (↑) |
|---|---|---|---|---|---|---|---|
| | | Dice (↑) | Overlap (↑) | Overreach (↓) | VS (↑) | IS (↓) | |
| AF_L | 0.5724 ± 0.1948 | 0.3913 ± 0.0866 | 0.3499 ± 0.1043 | 0.4142 ± 0.1727 | 0.5853 ± 0.1914 | 0.4147 ± 0.1914 | 0.2762 ± 0.0959 |
| AF_R | 0.6474 ± 0.1659 | 0.3430 ± 0.0902 | 0.3785 ± 0.1442 | 0.8125 ± 0.5997 | 0.6952 ± 0.1769 | 0.3048 ± 0.1769 | 0.2979 ± 0.1215 |
| CG_L | 0.6135 ± 0.2226 | 0.4357 ± 0.0808 | 0.5109 ± 0.0899 | 0.8959 ± 0.5376 | 0.5451 ± 0.2249 | 0.4549 ± 0.2249 | 0.4078 ± 0.0794 |
| CG_R | 0.7085 ± 0.1142 | 0.4719 ± 0.0773 | 0.5252 ± 0.0821 | 0.7566 ± 0.4827 | 0.5784 ± 0.1563 | 0.4216 ± 0.1563 | 0.4143 ± 0.0653 |
| FPT_L | 0.6786 ± 0.1733 | 0.4165 ± 0.0940 | 0.3099 ± 0.0874 | 0.1680 ± 0.1442 | 0.7904 ± 0.1814 | 0.2096 ± 0.1814 | 0.2668 ± 0.0779 |
| FPT_R | 0.6602 ± 0.2044 | 0.4193 ± 0.0777 | 0.3131 ± 0.0827 | 0.1724 ± 0.1677 | 0.7976 ± 0.2140 | 0.2024 ± 0.2140 | 0.2713 ± 0.0727 |
| IFOF_L | 0.4519 ± 0.1644 | 0.3111 ± 0.0914 | 0.2176 ± 0.0728 | 0.1643 ± 0.0728 | 0.7510 ± 0.1899 | 0.2490 ± 0.1899 | 0.1770 ± 0.0651 |
| IFOF_R | 0.5312 ± 0.2162 | 0.3354 ± 0.0853 | 0.2372 ± 0.0742 | 0.1509 ± 0.0656 | 0.7783 ± 0.1998 | 0.2217 ± 0.1998 | 0.1910 ± 0.0656 |
| ILF_L | 0.6701 ± 0.1553 | 0.2837 ± 0.0808 | 0.1901 ± 0.0768 | 0.1192 ± 0.1007 | 0.8825 ± 0.1192 | 0.1175 ± 0.1192 | 0.1577 ± 0.0663 |
| ILF_R | 0.6213 ± 0.1360 | 0.2893 ± 0.0620 | 0.1889 ± 0.0484 | 0.1090 ± 0.0743 | 0.8387 ± 0.1393 | 0.1613 ± 0.1393 | 0.1527 ± 0.0440 |
| MCP | 0.7205 ± 0.1817 | 0.5069 ± 0.0865 | 0.4824 ± 0.1200 | 0.4119 ± 0.2317 | 0.7570 ± 0.1241 | 0.2430 ± 0.1241 | 0.3967 ± 0.1091 |
| MdLF_L | 0.5075 ± 0.1888 | 0.3049 ± 0.0999 | 0.4034 ± 0.0909 | 1.9252 ± 2.5943 | 0.3182 ± 0.1912 | 0.6818 ± 0.1912 | 0.3036 ± 0.0685 |
| MdLF_R | 0.5203 ± 0.1581 | 0.2751 ± 0.1089 | 0.3404 ± 0.1023 | 2.0328 ± 3.1140 | 0.3122 ± 0.1565 | 0.6878 ± 0.1565 | 0.2611 ± 0.0874 |
| POPT_L | 0.6542 ± 0.1308 | 0.3936 ± 0.0647 | 0.3128 ± 0.0756 | 0.2657 ± 0.1733 | 0.5605 ± 0.1585 | 0.4395 ± 0.1585 | 0.2701 ± 0.0648 |
| POPT_R | 0.6414 ± 0.1061 | 0.4076 ± 0.0702 | 0.3371 ± 0.0903 | 0.3063 ± 0.2553 | 0.5253 ± 0.1330 | 0.4747 ± 0.1330 | 0.2923 ± 0.0800 |
| UF_L | 0.3124 ± 0.2656 | 0.2592 ± 0.1110 | 0.2161 ± 0.0994 | 0.5523 ± 0.5707 | 0.2976 ± 0.2775 | 0.7024 ± 0.2775 | 0.1715 ± 0.0836 |
| UF_R | 0.3555 ± 0.2821 | 0.3054 ± 0.0943 | 0.2613 ± 0.1060 | 0.5263 ± 0.6797 | 0.2822 ± 0.2667 | 0.7178 ± 0.2667 | 0.1968 ± 0.0874 |

**Tab. S7 | Geometry and connectivity of bundles generated with ATM.** Absolute percent differences are presented as mean ± standard deviation. A dash '−' indicates an absent bundle.

| Bundles | Geometry APD (↓) | | | | Pearson (↑) | Connectivity APD (↓) | | | |
|---|---|---|---|---|---|---|---|---|---|
| | Mean Length | Span | Volume | Surface Area | | Density | CPL | Efficiency | Modularity |
| AF_L | 16.3782 ± 6.1705 | 11.9295 ± 9.4784 | 31.7351 ± 21.8256 | 48.1792 ± 28.5575 | 0.4791 ± 0.1895 | 71.5249 ± 43.9653 | 113.4541 ± 50.0194 | 147.1505 ± 30.8938 | 97.5305 ± 54.6368 |
| AF_R | 17.1918 ± 6.3730 | 10.4354 ± 6.7693 | 53.6095 ± 34.0503 | 64.8241 ± 34.2957 | 0.4818 ± 0.1450 | 84.7245 ± 39.3843 | 103.6713 ± 49.5844 | 162.7604 ± 19.4216 | 104.1599 ± 57.3431 |
| CC_Fr_1 | 13.0568 ± 4.7349 | 12.0143 ± 9.6003 | 43.5380 ± 27.7355 | 30.6197 ± 18.7808 | 0.5115 ± 0.2119 | 77.4166 ± 75.8007 | 111.4594 ± 65.6328 | 116.9450 ± 63.9403 | 117.8747 ± 90.3340 |
| CC_Fr_2 | 18.9026 ± 4.6200 | 23.4412 ± 9.4057 | 71.5429 ± 31.9900 | 33.3150 ± 19.6487 | 0.5941 ± 0.1848 | 53.6970 ± 44.9040 | 118.2380 ± 52.1721 | 120.4965 ± 46.8038 | 154.7775 ± 65.3555 |
| CC_Oc | 11.0030 ± 6.1956 | 10.9535 ± 7.9614 | 21.7280 ± 16.1675 | 29.7788 ± 16.9099 | 0.5395 ± 0.1194 | 26.4474 ± 18.3870 | 59.1651 ± 31.3135 | 119.3264 ± 24.0460 | 161.2685 ± 67.0146 |
| CC_Pa | 12.0806 ± 4.7389 | 17.5031 ± 10.8307 | 44.8484 ± 24.8365 | 23.9097 ± 18.3151 | 0.5634 ± 0.0817 | 22.6523 ± 13.1722 | 60.9603 ± 17.3834 | 91.3646 ± 23.7189 | 161.6402 ± 70.5580 |
| CC_Pr_Po | 14.5432 ± 6.2803 | 20.1826 ± 10.0380 | 62.9048 ± 18.3201 | 20.0317 ± 10.2214 | 0.5122 ± 0.1061 | 28.3413 ± 20.7427 | 68.2592 ± 32.2108 | 91.9094 ± 23.6135 | 179.1153 ± 45.5032 |
| CG_L | 11.7686 ± 8.1369 | 7.8983 ± 6.0824 | 43.0467 ± 31.4850 | 50.6015 ± 25.2450 | 0.5361 ± 0.1424 | 63.9521 ± 37.4996 | 80.8836 ± 57.6416 | 149.4564 ± 29.5670 | 105.5089 ± 84.1536 |
| CG_R | 14.4626 ± 5.5075 | 9.1331 ± 5.2334 | 42.4491 ± 31.5814 | 50.7069 ± 25.2823 | 0.4689 ± 0.1448 | 70.4776 ± 43.0928 | 91.8669 ± 52.1589 | 165.4556 ± 18.8717 | 119.0609 ± 76.6755 |
| FAT_L | 22.4657 ± 5.0508 | 16.0017 ± 5.7259 | 39.1771 ± 18.4583 | 33.0898 ± 21.5166 | 0.5763 ± 0.1146 | 36.7774 ± 45.0994 | 137.6900 ± 46.0749 | 90.8868 ± 38.1938 | 149.8390 ± 77.2425 |
| FAT_R | 24.5213 ± 5.0085 | 18.3513 ± 6.1703 | 33.3269 ± 21.8262 | 33.5119 ± 20.0700 | 0.6050 ± 0.1024 | 53.7432 ± 51.0949 | 135.9431 ± 51.0177 | 117.7373 ± 34.7561 | 138.4228 ± 71.0034 |
| FPT_L | 12.2648 ± 6.0247 | 8.7889 ± 6.0314 | 32.5808 ± 21.2695 | 29.1909 ± 25.6331 | 0.3836 ± 0.2600 | 96.7651 ± 51.7182 | 126.7067 ± 48.9923 | 162.8325 ± 34.8590 | 157.5421 ± 54.7153 |
| FPT_R | 12.6060 ± 6.6058 | 10.2867 ± 6.3087 | 36.9889 ± 19.8639 | 25.3915 ± 26.2654 | 0.3577 ± 0.2355 | 84.2431 ± 46.7789 | 138.3907 ± 51.2070 | 158.0912 ± 34.4026 | 108.2244 ± 66.8066 |
| IFOF_L | 8.1415 ± 5.1921 | 6.3258 ± 3.8312 | 22.8766 ± 14.6966 | 27.1602 ± 15.1998 | 0.5190 ± 0.1150 | 41.2266 ± 44.6142 | 90.4319 ± 47.4766 | 136.7237 ± 30.9487 | 99.6624 ± 59.8478 |
| IFOF_R | 6.4569 ± 3.9587 | 5.6194 ± 3.6281 | 19.9552 ± 11.0831 | 19.6940 ± 11.2226 | 0.5350 ± 0.1568 | 42.6251 ± 25.6825 | 76.4801 ± 52.7638 | 133.4644 ± 34.3376 | 119.1448 ± 53.8623 |
| ILF_L | 13.4611 ± 6.1576 | 8.9987 ± 5.0363 | 36.7725 ± 16.7823 | 36.6135 ± 28.1284 | 0.3397 ± 0.1190 | 49.4572 ± 44.9010 | 90.5238 ± 48.4326 | 130.3446 ± 36.1894 | 109.2040 ± 67.3970 |
| ILF_R | 15.7252 ± 10.8096 | 10.4027 ± 7.7099 | 34.3426 ± 19.5810 | 41.6718 ± 34.7760 | 0.3617 ± 0.1705 | 35.0685 ± 31.5976 | 75.5435 ± 33.9021 | 116.6163 ± 34.5262 | 122.6967 ± 75.3571 |
| MCP | 12.7934 ± 10.3452 | 18.9207 ± 15.4606 | 41.3428 ± 30.9230 | 58.8321 ± 36.0299 | - | - | - | - | - |
| MdLF_L | 9.5037 ± 8.1335 | 8.1337 ± 6.3170 | 41.3472 ± 38.9247 | 67.5858 ± 31.4190 | 0.4828 ± 0.1114 | 58.2989 ± 37.5180 | 74.9776 ± 34.2174 | 136.8213 ± 23.7961 | 96.1600 ± 67.7045 |
| MdLF_R | 13.5569 ± 6.2466 | 8.1571 ± 7.1910 | 34.5415 ± 31.7768 | 61.2891 ± 28.3313 | 0.4888 ± 0.1381 | 55.4073 ± 38.5082 | 73.2844 ± 24.6850 | 136.8928 ± 23.3823 | 112.0057 ± 72.1071 |
| OR_ML_L | 8.6944 ± 5.5935 | 8.0702 ± 5.2010 | 36.0904 ± 27.1998 | 53.1422 ± 24.7711 | 0.5025 ± 0.1239 | 43.8691 ± 36.3039 | 73.8708 ± 28.7495 | 130.0176 ± 27.2128 | 93.7145 ± 64.2978 |
| OR_ML_R | 6.5757 ± 5.3304 | 9.1249 ± 6.4066 | 59.6100 ± 30.9532 | 69.7776 ± 25.9030 | 0.3820 ± 0.1415 | 42.5585 ± 37.0258 | 72.2283 ± 30.7377 | 129.8328 ± 25.5756 | 62.2036 ± 59.0656 |
| POPT_L | 11.8635 ± 5.1048 | 7.9826 ± 4.4917 | 27.9131 ± 16.9057 | 24.9565 ± 15.8569 | 0.3151 ± 0.1891 | 56.1726 ± 42.1884 | 88.7277 ± 57.0255 | 127.6431 ± 47.9636 | 83.6436 ± 73.2118 |
| POPT_R | 12.8781 ± 4.3365 | 10.5900 ± 4.6060 | 30.6801 ± 19.0879 | 29.4745 ± 20.2749 | 0.2805 ± 0.1576 | 58.3800 ± 56.2500 | 115.2927 ± 47.8857 | 149.0068 ± 27.3337 | 87.8550 ± 57.3309 |
| PYT_L | 12.5660 ± 6.8230 | 10.3589 ± 6.6500 | 52.9165 ± 16.2892 | 11.0647 ± 9.7582 | 0.3818 ± 0.2085 | 59.2035 ± 51.5608 | 112.3338 ± 57.5605 | 118.9285 ± 47.3877 | 111.2434 ± 69.3786 |
| PYT_R | 10.5912 ± 6.6160 | 8.9615 ± 6.4546 | 51.4753 ± 13.1687 | 13.7194 ± 7.5828 | 0.3414 ± 0.2046 | 50.9705 ± 39.8734 | 120.1050 ± 45.0122 | 133.3500 ± 33.4591 | 83.5500 ± 63.0634 |
| SLF_L | 16.0281 ± 8.4210 | 10.6847 ± 8.2703 | 48.6050 ± 29.7418 | 69.8918 ± 27.8327 | 0.3965 ± 0.1766 | 113.5328 ± 53.9868 | 132.9065 ± 50.1194 | 175.9793 ± 16.6950 | 126.1223 ± 68.6660 |
| SLF_R | 14.1764 ± 7.4493 | 7.7023 ± 6.8997 | 43.9492 ± 29.5444 | 59.8345 ± 32.0203 | 0.4961 ± 0.1586 | 68.8449 ± 48.7939 | 102.3647 ± 44.0100 | 154.9991 ± 22.3582 | 108.8233 ± 66.3844 |
| UF_L | 9.5315 ± 7.0647 | 10.9449 ± 8.8163 | 48.5047 ± 25.9682 | 66.7306 ± 30.4244 | 0.5044 ± 0.1572 | 91.2972 ± 62.8138 | 68.1320 ± 59.6366 | 133.9668 ± 51.3670 | 106.1518 ± 83.9562 |
| UF_R | 12.1006 ± 6.5600 | 10.9832 ± 7.3216 | 30.0130 ± 26.1309 | 46.0246 ± 31.3479 | 0.4548 ± 0.1819 | 63.6916 ± 67.1990 | 70.4086 ± 44.0482 | 105.7981 ± 56.7679 | 132.1564 ± 74.7255 |

**Tab. S8 | Geometry and connectivity of bundles generated with ATM-Population.** Absolute percent differences are presented as mean $\pm$ standard deviation. A dash '$-$' indicates an absent bundle.

| Bundles | Geometry APD (↓) | | | | Pearson (↑) | Connectivity APD (↓) | | | |
|---|---|---|---|---|---|---|---|---|---|
| | Mean Length | Span | Volume | Surface Area | | Density | CPL | Efficiency | Modularity |
| AF_L | 15.1903 ± 7.4526 | 14.6485 ± 11.5812 | 35.9787 ± 22.1739 | 43.4696 ± 32.6299 | 0.3844 ± 0.1669 | 67.3214 ± 46.1348 | 113.9775 ± 52.1029 | 143.0450 ± 32.6974 | 87.3707 ± 64.9730 |
| AF_R | 14.9581 ± 8.4364 | 12.7552 ± 6.8572 | 65.1228 ± 43.3370 | 68.9540 ± 39.7595 | 0.4326 ± 0.1512 | 90.5512 ± 40.3479 | 95.0792 ± 54.3016 | 160.9776 ± 26.3262 | 104.5312 ± 53.8798 |
| CC_Fr_1 | 15.8534 ± 7.9409 | 17.8287 ± 15.5669 | 65.6969 ± 49.4697 | 44.6863 ± 45.4319 | 0.5193 ± 0.1558 | 85.7088 ± 72.6479 | 135.3560 ± 59.4086 | 110.8318 ± 66.5151 | 97.0505 ± 98.3693 |
| CC_Fr_2 | 12.7780 ± 9.5932 | 17.6194 ± 20.0278 | 84.8404 ± 46.4618 | 48.6147 ± 39.9566 | 0.6440 ± 0.1734 | 60.6623 ± 58.4476 | 99.9237 ± 52.4488 | 120.8715 ± 51.8214 | 147.7213 ± 70.1557 |
| CC_Oc | 15.7544 ± 7.5255 | 13.2561 ± 9.9175 | 42.8328 ± 21.7515 | 20.8672 ± 14.5946 | 0.4249 ± 0.1506 | 32.7311 ± 18.7321 | 59.9813 ± 31.8156 | 107.1783 ± 27.0847 | 161.8904 ± 65.3469 |
| CC_Pa | 17.3933 ± 9.1248 | 16.3454 ± 17.5016 | 67.6609 ± 24.2470 | 27.6428 ± 18.0493 | 0.4745 ± 0.1174 | 29.8949 ± 18.3837 | 63.4011 ± 21.1566 | 86.5678 ± 21.7243 | 166.8034 ± 62.7219 |
| CC_Pr_Po | 18.7061 ± 11.2731 | 17.3071 ± 15.8513 | 81.3522 ± 25.8039 | 30.0435 ± 19.3799 | 0.4794 ± 0.1305 | 36.1153 ± 24.9189 | 74.5614 ± 28.3515 | 81.3426 ± 22.1037 | 164.6621 ± 67.0292 |
| CG_L | 15.8380 ± 10.0238 | 11.5765 ± 7.6402 | 35.5273 ± 27.2557 | 40.3387 ± 24.7498 | 0.4735 ± 0.1800 | 55.3990 ± 44.8968 | 94.4838 ± 56.4260 | 137.0613 ± 35.8240 | 104.3670 ± 88.3807 |
| CG_R | 16.2125 ± 7.7357 | 12.2955 ± 7.4877 | 32.7588 ± 30.4698 | 44.2456 ± 25.7716 | 0.4075 ± 0.1931 | 69.2777 ± 45.5215 | 110.7252 ± 46.9127 | 158.4514 ± 24.6276 | 135.7073 ± 76.8905 |
| FAT_L | 19.0706 ± 8.8019 | 18.9685 ± 8.8322 | 66.9717 ± 42.2065 | 31.3912 ± 39.8544 | 0.5329 ± 0.1505 | 28.4296 ± 23.1792 | 121.1555 ± 49.5196 | 78.0864 ± 36.0665 | 144.6450 ± 79.1146 |
| FAT_R | 17.7398 ± 11.7693 | 16.4434 ± 11.1929 | 62.4829 ± 45.6368 | 37.0495 ± 34.5284 | 0.5468 ± 0.1407 | 48.0135 ± 38.4612 | 111.2467 ± 53.1970 | 106.0854 ± 39.8015 | 133.3885 ± 77.2468 |
| FPT_L | 15.9829 ± 7.1604 | 16.9720 ± 6.7668 | 31.0011 ± 20.7129 | 30.4112 ± 23.2669 | 0.3394 ± 0.2522 | 90.4185 ± 51.1181 | 140.1449 ± 38.4490 | 162.8316 ± 23.5165 | 144.8132 ± 66.1390 |
| FPT_R | 15.1898 ± 7.8461 | 18.0467 ± 7.5642 | 36.2902 ± 19.5963 | 27.9517 ± 23.8032 | 0.3533 ± 0.2401 | 93.4268 ± 49.2626 | 154.6930 ± 39.0232 | 166.9447 ± 22.4762 | 116.1346 ± 60.8716 |
| IFOF_L | 9.5513 ± 6.2685 | 7.0166 ± 5.1567 | 31.6527 ± 20.8527 | 14.0585 ± 10.0268 | 0.4649 ± 0.1078 | 49.2110 ± 50.8075 | 105.3128 ± 45.2055 | 131.4743 ± 33.2149 | 123.8310 ± 58.5869 |
| IFOF_R | 6.9876 ± 4.3555 | 7.5880 ± 5.4508 | 22.7739 ± 15.3100 | 19.0003 ± 9.2328 | 0.4848 ± 0.1854 | 51.1656 ± 34.7996 | 86.0917 ± 55.8142 | 127.0845 ± 36.4778 | 118.8417 ± 59.7063 |
| ILF_L | 14.4210 ± 6.3183 | 10.5947 ± 5.5977 | 43.9348 ± 18.0770 | 20.4695 ± 21.6554 | 0.2617 ± 0.1143 | 50.0603 ± 48.5389 | 109.1577 ± 48.4228 | 130.5510 ± 36.0233 | 118.0520 ± 65.4610 |
| ILF_R | 11.2728 ± 5.4289 | 7.9269 ± 5.3997 | 33.9551 ± 18.9097 | 19.3101 ± 16.6968 | 0.3127 ± 0.1335 | 26.8452 ± 26.0498 | 95.8546 ± 37.9261 | 124.1816 ± 21.0852 | 119.7859 ± 77.0494 |
| MCP | 14.9683 ± 11.2561 | 17.8836 ± 14.9682 | 32.5357 ± 23.0735 | 44.0487 ± 36.1709 | - | - | - | - | - |
| MdLF_L | 19.8016 ± 10.0455 | 17.9470 ± 8.6171 | 55.5269 ± 48.8826 | 50.5459 ± 32.5481 | 0.4016 ± 0.1465 | 63.3060 ± 41.2535 | 88.6555 ± 38.8299 | 135.4596 ± 28.6010 | 97.6667 ± 68.5319 |
| MdLF_R | 19.0719 ± 9.7041 | 16.4447 ± 10.7364 | 55.1219 ± 51.5649 | 55.8986 ± 41.4869 | 0.4476 ± 0.1573 | 65.5988 ± 45.3818 | 78.6727 ± 27.0928 | 138.5169 ± 31.4171 | 117.2157 ± 67.0735 |
| OR_ML_L | 9.9549 ± 5.8899 | 9.2682 ± 5.6871 | 33.4464 ± 24.4813 | 33.2180 ± 24.2768 | 0.4035 ± 0.1988 | 44.1277 ± 35.3642 | 70.0507 ± 34.3818 | 116.7809 ± 35.4810 | 112.1680 ± 63.4526 |
| OR_ML_R | 6.4981 ± 6.2865 | 11.4850 ± 6.7212 | 57.7529 ± 43.6939 | 54.3059 ± 37.4020 | 0.2694 ± 0.1658 | 43.6361 ± 45.2740 | 67.2715 ± 35.4010 | 118.9910 ± 37.4315 | 68.3848 ± 60.4036 |
| POPT_L | 20.9558 ± 8.3149 | 22.6443 ± 7.0660 | 42.3865 ± 37.4088 | 25.3714 ± 32.0418 | 0.2751 ± 0.1665 | 68.0210 ± 57.2305 | 119.9637 ± 44.6489 | 149.1643 ± 29.2007 | 98.2046 ± 75.6435 |
| POPT_R | 18.0135 ± 7.0777 | 23.1338 ± 7.7626 | 42.9306 ± 36.2515 | 27.7317 ± 31.1224 | 0.2708 ± 0.1782 | 71.9133 ± 59.4201 | 131.8678 ± 50.0063 | 149.8203 ± 31.8575 | 97.7578 ± 69.3978 |
| PYT_L | 19.6281 ± 10.1564 | 23.6654 ± 9.0217 | 63.4540 ± 31.9547 | 21.2143 ± 30.5960 | 0.2747 ± 0.1685 | 60.8503 ± 54.6519 | 138.8670 ± 44.2535 | 132.3403 ± 39.4252 | 126.1570 ± 65.5734 |
| PYT_R | 13.6734 ± 8.1576 | 17.4876 ± 8.8137 | 47.8747 ± 14.9153 | 14.6446 ± 9.9401 | 0.3635 ± 0.2185 | 52.6821 ± 54.5900 | 133.0168 ± 47.4381 | 143.4494 ± 27.6561 | 95.3623 ± 66.7776 |
| SLF_L | 11.9386 ± 8.5120 | 12.3235 ± 8.6830 | 73.4610 ± 50.5316 | 79.9947 ± 40.0558 | 0.3565 ± 0.1669 | 95.2981 ± 58.8599 | 92.1867 ± 63.8907 | 161.9181 ± 38.4239 | 97.0878 ± 71.7606 |
| SLF_R | 11.8938 ± 6.9515 | 8.5291 ± 8.4121 | 53.3716 ± 36.9150 | 52.5401 ± 37.7580 | 0.4816 ± 0.1593 | 63.9322 ± 51.4226 | 92.9967 ± 55.1508 | 148.2471 ± 26.3525 | 113.8902 ± 72.1118 |
| UF_L | 23.6261 ± 10.5987 | 13.6764 ± 9.6093 | 69.5496 ± 47.5604 | 61.6631 ± 33.2409 | 0.3876 ± 0.1714 | 96.6155 ± 65.2664 | 120.5505 ± 52.3327 | 148.1419 ± 38.7502 | 143.3888 ± 78.5278 |
| UF_R | 17.9449 ± 6.2685 | 13.6795 ± 10.1713 | 52.6053 ± 46.9771 | 52.8957 ± 35.9998 | 0.3329 ± 0.1779 | 76.0534 ± 67.1183 | 106.8625 ± 54.9917 | 133.0591 ± 46.0476 | 140.4326 ± 83.3807 |

**Tab. S9 | Geometry and connectivity of bundles generated from MRtrix with BundleSeg.** Absolute percent differences are presented as mean ± standard deviation. A dash '−' indicates an absent bundle.

| Bundles | Geometry APD (↓) | | | | Pearson (↑) | Connectivity APD (↓) | | | |
|---|---|---|---|---|---|---|---|---|---|
| | Mean Length | Span | Volume | Surface Area | | Density | CPL | Efficiency | Modularity |
| AF_L | 28.8638 ± 6.2594 | 13.8137 ± 9.3045 | 87.9822 ± 50.4303 | 68.7963 ± 44.9017 | 0.4425 ± 0.2088 | 56.2553 ± 38.0790 | 115.9907 ± 50.9674 | 92.1982 ± 51.5560 | 105.2410 ± 73.4941 |
| AF_R | 29.3859 ± 6.8291 | 17.1100 ± 13.3408 | 73.9471 ± 53.0796 | 58.2679 ± 44.7810 | 0.4297 ± 0.2057 | 44.3435 ± 37.8135 | 120.6064 ± 49.0598 | 115.1491 ± 43.6608 | 97.9310 ± 70.8317 |
| CC_Fr_1 | 5.7839 ± 3.7027 | 43.7240 ± 20.1852 | 78.7807 ± 55.1777 | 54.2505 ± 54.1674 | 0.0850 ± 0.1963 | 112.9959 ± 92.1093 | 32.3852 ± 55.1185 | 110.0034 ± 92.7009 | 95.4459 ± 99.6944 |
| CC_Fr_2 | 2.7875 ± 2.6557 | 77.9988 ± 20.6141 | 44.1374 ± 44.1312 | 45.1637 ± 37.7295 | 0.4173 ± 0.2821 | 84.4714 ± 59.0603 | 78.1880 ± 57.8595 | 76.5083 ± 62.6640 | 133.0280 ± 82.0938 |
| CC_Oc | 35.7338 ± 4.4085 | 32.8104 ± 17.8196 | 177.0229 ± 19.2431 | 156.8045 ± 23.9637 | 0.2808 ± 0.2287 | 168.4303 ± 26.6293 | 75.6919 ± 47.6330 | 121.8974 ± 63.8347 | 104.2370 ± 97.1341 |
| CC_Pa | 14.3383 ± 4.2941 | 32.4736 ± 13.3728 | 77.7381 ± 41.4743 | 48.1654 ± 34.0950 | 0.4698 ± 0.1117 | 38.9257 ± 40.9147 | 60.3066 ± 21.8955 | 73.7371 ± 33.9708 | 169.2051 ± 59.6969 |
| CC_Pr_Po | 12.9030 ± 4.1850 | 67.7001 ± 17.2934 | 29.1074 ± 35.7807 | 26.2781 ± 27.9477 | 0.4167 ± 0.1398 | 39.0192 ± 40.9505 | 46.9349 ± 31.5004 | 88.4911 ± 23.5332 | 164.8780 ± 67.2973 |
| CG_L | 17.8938 ± 13.7226 | 14.0711 ± 11.9538 | 69.1726 ± 23.1860 | 75.6729 ± 21.5253 | 0.4303 ± 0.1663 | 110.8540 ± 48.8508 | 107.3108 ± 46.5270 | 172.8764 ± 23.9367 | 136.6753 ± 65.7551 |
| CG_R | 16.7167 ± 17.4993 | 12.2994 ± 14.1988 | 80.6849 ± 28.6274 | 85.7612 ± 24.1933 | 0.3165 ± 0.1820 | 125.1664 ± 42.2362 | 113.3683 ± 50.5499 | 179.1848 ± 19.0918 | 157.8087 ± 63.9693 |
| FAT_L | 5.2375 ± 3.7237 | 8.1201 ± 5.1925 | 41.3541 ± 42.6404 | 30.0200 ± 37.0744 | 0.3467 ± 0.2989 | 117.9954 ± 64.1643 | 112.7302 ± 55.6808 | 82.4727 ± 86.1731 | 78.5563 ± 94.9835 |
| FAT_R | 5.3931 ± 4.0602 | 7.0737 ± 4.7337 | 33.0693 ± 44.4301 | 31.7190 ± 41.1119 | 0.3900 ± 0.2727 | 99.1829 ± 67.7925 | 101.1498 ± 59.5998 | 91.7444 ± 65.1141 | 126.3478 ± 91.3138 |
| FPT_L | 21.8017 ± 8.4264 | 18.0467 ± 7.5873 | 129.1507 ± 35.1542 | 100.6918 ± 37.1380 | 0.2330 ± 0.2834 | 110.6044 ± 64.7949 | 73.4742 ± 58.0104 | 97.6367 ± 67.7734 | 120.6616 ± 94.9560 |
| FPT_R | 21.1294 ± 7.9463 | 17.3374 ± 7.5307 | 117.0144 ± 47.3696 | 92.9250 ± 45.1590 | 0.2453 ± 0.2493 | 99.7060 ± 60.4912 | 98.5897 ± 62.4057 | 81.1510 ± 70.6249 | 137.8028 ± 87.3037 |
| IFOF_L | 41.6731 ± 8.0817 | 33.0382 ± 8.3303 | 146.0696 ± 24.1083 | 137.6828 ± 21.9983 | 0.2132 ± 0.1760 | 120.9023 ± 38.4317 | 127.9149 ± 56.9797 | 61.3328 ± 53.1011 | 161.9909 ± 59.4537 |
| IFOF_R | 37.8219 ± 8.1011 | 28.2536 ± 8.8492 | 143.9668 ± 23.8343 | 139.3631 ± 19.7904 | 0.2836 ± 0.1989 | 92.4370 ± 60.3363 | 114.7265 ± 55.7990 | 87.1881 ± 41.8378 | 161.8784 ± 48.8645 |
| ILF_L | 15.8867 ± 6.4041 | 11.8636 ± 6.1887 | 31.7209 ± 33.2591 | 30.6702 ± 31.6747 | 0.3473 ± 0.1449 | 39.3282 ± 43.0886 | 80.9569 ± 35.1834 | 108.2052 ± 40.1258 | 122.3668 ± 73.5356 |
| ILF_R | 14.7783 ± 6.1654 | 11.5663 ± 5.0512 | 33.2335 ± 32.3118 | 32.5480 ± 31.8193 | 0.3683 ± 0.1289 | 26.4232 ± 28.0865 | 70.8102 ± 34.0117 | 103.4886 ± 31.6855 | 142.5396 ± 62.3275 |
| MCP | 28.2028 ± 10.7673 | 22.4281 ± 23.0801 | 146.4465 ± 61.9727 | 125.2056 ± 54.0225 | - | - | - | - | - |
| MdLF_L | 8.7013 ± 6.9419 | 8.6567 ± 7.7469 | 51.6726 ± 33.2609 | 39.0602 ± 29.6036 | 0.4715 ± 0.1562 | 42.1907 ± 33.0752 | 85.1519 ± 36.7559 | 105.0234 ± 37.3388 | 104.6511 ± 66.9734 |
| MdLF_R | 6.3598 ± 4.8500 | 6.6968 ± 4.8696 | 57.4872 ± 46.5521 | 47.8694 ± 39.0933 | 0.4932 ± 0.1539 | 53.7005 ± 37.4463 | 68.7817 ± 20.6471 | 118.9414 ± 40.5299 | 116.3487 ± 70.6585 |
| OR_ML_L | 5.0850 ± 4.4191 | 5.8998 ± 4.8042 | 53.9135 ± 32.0633 | 31.1518 ± 22.3581 | 0.3963 ± 0.1263 | 54.2445 ± 21.1140 | 74.9794 ± 30.5771 | 74.2082 ± 36.6191 | 85.3263 ± 66.2089 |
| OR_ML_R | 6.1623 ± 7.3072 | 8.2874 ± 7.1831 | 63.4450 ± 40.7029 | 37.5376 ± 31.3197 | 0.2923 ± 0.1551 | 63.8084 ± 39.9575 | 77.1344 ± 34.6310 | 79.4531 ± 48.7820 | 96.9684 ± 57.8442 |
| POPT_L | 19.0844 ± 5.3474 | 19.3754 ± 6.5343 | 120.3868 ± 40.6702 | 94.4559 ± 34.5649 | 0.3025 ± 0.2467 | 121.6281 ± 54.9378 | 73.1873 ± 55.7223 | 82.3611 ± 69.6629 | 105.3239 ± 77.2181 |
| POPT_R | 18.8785 ± 4.7430 | 20.5395 ± 4.8500 | 115.9403 ± 44.8431 | 93.6892 ± 34.9535 | 0.3039 ± 0.2123 | 99.3844 ± 62.0769 | 73.5053 ± 60.3639 | 77.7852 ± 61.7575 | 115.3905 ± 78.0092 |
| PYT_L | 17.0738 ± 7.7422 | 15.1706 ± 8.3321 | 88.5464 ± 36.6111 | 72.6984 ± 34.1138 | 0.3723 ± 0.2440 | 117.2104 ± 60.4074 | 82.2833 ± 64.3473 | 91.9005 ± 66.2040 | 108.1071 ± 92.0837 |
| PYT_R | 17.5932 ± 7.2573 | 16.5858 ± 7.6242 | 91.4248 ± 27.7694 | 74.7841 ± 24.3140 | 0.3465 ± 0.2431 | 115.0554 ± 56.7531 | 77.4424 ± 64.5653 | 83.1131 ± 65.8201 | 115.0921 ± 75.0838 |
| SLF_L | 26.7878 ± 36.5661 | 24.5800 ± 36.9272 | 75.9983 ± 53.6756 | 62.5151 ± 49.2701 | 0.3507 ± 0.2348 | 101.9027 ± 56.8448 | 101.5612 ± 59.8188 | 159.8084 ± 44.2256 | 103.7727 ± 71.2417 |
| SLF_R | 23.1124 ± 36.9801 | 18.8120 ± 37.7264 | 64.1922 ± 56.0286 | 57.3220 ± 48.9318 | 0.5139 ± 0.1740 | 67.5001 ± 55.2840 | 110.9925 ± 52.5646 | 147.4819 ± 32.0273 | 98.8109 ± 73.5318 |
| UF_L | 7.9091 ± 9.8992 | 15.3006 ± 12.0420 | 76.9889 ± 45.3026 | 56.9821 ± 41.7263 | 0.2605 ± 0.2369 | 57.4282 ± 70.8179 | 56.1935 ± 51.6789 | 97.5877 ± 65.2284 | 102.2076 ± 84.5329 |
| UF_R | 6.3512 ± 6.0436 | 12.6905 ± 9.1482 | 52.6721 ± 43.3370 | 47.1954 ± 37.9493 | 0.3177 ± 0.2492 | 59.0160 ± 68.4713 | 22.7814 ± 18.9467 | 93.2299 ± 62.8052 | 88.8137 ± 77.5006 |

**Tab. S10 | Geometry and connectivity of bundles generated with SCIL WM atlas warping.** Absolute percent differences are presented as mean ± standard deviation. A dash '−' indicates an absent bundle.

| Bundles | Geometry APD (↓) | | | | Pearson (↑) | Connectivity APD (↓) | | | |
|---|---|---|---|---|---|---|---|---|---|
| | Mean Length | Span | Volume | Surface Area | | Density | CPL | Efficiency | Modularity |
| AF_L | 6.8671 ± 5.1370 | 12.9578 ± 9.6855 | 80.4248 ± 31.5824 | 65.8603 ± 29.4192 | 0.5117 ± 0.1435 | 65.3658 ± 54.7625 | 72.8656 ± 45.7535 | 139.0153 ± 37.6715 | 113.2092 ± 59.4608 |
| AF_R | 6.2998 ± 5.2393 | 11.7393 ± 6.2468 | 110.7677 ± 43.0565 | 87.1237 ± 37.6548 | 0.4812 ± 0.1484 | 87.0591 ± 34.9837 | 59.9066 ± 41.0402 | 163.0339 ± 17.7238 | 105.9860 ± 60.0702 |
| CC_Fr_1 | 15.3850 ± 5.3386 | 69.1439 ± 14.0994 | 52.5735 ± 29.4224 | 28.1593 ± 21.1176 | 0.2810 ± 0.0934 | 120.0259 ± 91.4889 | 36.0125 ± 56.2544 | 113.0662 ± 92.8543 | 86.7505 ± 98.9151 |
| CC_Fr_2 | 6.7779 ± 4.2808 | 79.2540 ± 17.3526 | 34.3718 ± 26.8075 | 34.7595 ± 18.6882 | 0.3184 ± 0.3133 | 93.9333 ± 54.4729 | 85.7562 ± 61.1966 | 57.1404 ± 59.0518 | 125.5198 ± 86.6299 |
| CC_Oc | 5.8269 ± 4.1057 | 26.1639 ± 11.7008 | 31.4334 ± 19.6175 | 34.0720 ± 15.8905 | 0.3774 ± 0.1972 | 55.4983 ± 42.8273 | 45.2238 ± 29.1175 | 75.1446 ± 36.5914 | 177.0998 ± 45.6462 |
| CC_Pa | 5.0622 ± 4.3203 | 69.1572 ± 13.2717 | 24.1413 ± 17.6771 | 39.7097 ± 23.2796 | 0.4106 ± 0.1169 | 43.7868 ± 16.2084 | 60.3170 ± 16.2961 | 73.3030 ± 24.1981 | 162.7212 ± 69.0877 |
| CC_Pr_Po | 7.2905 ± 3.6814 | 76.4518 ± 15.2194 | 20.2895 ± 10.3475 | 19.0045 ± 12.5899 | 0.4542 ± 0.1735 | 62.5391 ± 36.8505 | 52.4721 ± 36.9441 | 68.2240 ± 38.0621 | 165.5790 ± 65.4351 |
| CG_L | 9.3606 ± 7.1440 | 7.1445 ± 5.4248 | 65.1877 ± 32.5148 | 54.4920 ± 26.2758 | 0.3232 ± 0.2259 | 68.1916 ± 43.3152 | 73.1483 ± 58.6951 | 154.2046 ± 28.2237 | 120.0022 ± 74.5257 |
| CG_R | 7.5502 ± 4.7267 | 5.5390 ± 3.7249 | 70.8563 ± 28.5890 | 57.4128 ± 24.9052 | 0.1967 ± 0.1724 | 88.1679 ± 41.3946 | 79.2521 ± 61.9854 | 165.4979 ± 21.3220 | 169.8746 ± 51.8653 |
| FAT_L | 21.2143 ± 4.4211 | 27.4959 ± 4.9851 | 61.4267 ± 22.5394 | 40.8125 ± 15.7842 | 0.2899 ± 0.1970 | 147.8708 ± 65.1646 | 114.3985 ± 58.8828 | 131.7010 ± 76.2037 | 57.5395 ± 86.2745 |
| FAT_R | 18.0987 ± 6.1302 | 22.9826 ± 8.6943 | 47.2526 ± 22.3038 | 33.9039 ± 18.4856 | 0.3415 ± 0.1170 | 136.1283 ± 66.4507 | 111.4051 ± 59.7472 | 112.0758 ± 80.1887 | 117.4574 ± 98.2727 |
| FPT_L | 6.4878 ± 4.9733 | 5.1643 ± 5.2063 | 28.3680 ± 19.1025 | 24.6312 ± 15.9789 | 0.2449 ± 0.2240 | 140.3164 ± 72.0895 | 68.1552 ± 63.2570 | 132.9945 ± 69.4190 | 124.9392 ± 96.7776 |
| FPT_R | 5.3322 ± 4.2567 | 4.8863 ± 5.2827 | 30.3603 ± 20.3129 | 20.4533 ± 16.1621 | 0.2556 ± 0.2291 | 134.5413 ± 66.5576 | 70.5679 ± 61.7485 | 114.2451 ± 75.5957 | 149.5326 ± 82.3574 |
| IFOF_L | 5.5881 ± 4.1287 | 9.4986 ± 5.5994 | 28.1587 ± 19.2804 | 18.2430 ± 10.9580 | 0.4457 ± 0.1790 | 63.6361 ± 30.7008 | 46.5834 ± 40.0148 | 69.5504 ± 37.9557 | 139.7234 ± 64.4818 |
| IFOF_R | 7.7613 ± 5.6856 | 13.3749 ± 8.2729 | 24.2648 ± 15.8628 | 14.9966 ± 9.9397 | 0.5059 ± 0.1928 | 70.2414 ± 38.9035 | 56.6767 ± 46.5375 | 72.2860 ± 42.7081 | 131.4996 ± 70.1214 |
| ILF_L | 7.8329 ± 4.6974 | 5.2944 ± 3.6101 | 29.3315 ± 26.3861 | 23.8331 ± 17.6178 | 0.2635 ± 0.1469 | 54.2988 ± 49.7338 | 56.0230 ± 40.7227 | 80.4280 ± 48.8916 | 120.3754 ± 68.8230 |
| ILF_R | 5.1991 ± 4.1538 | 6.1044 ± 2.7182 | 24.9717 ± 21.1794 | 17.5200 ± 14.3292 | 0.2726 ± 0.1521 | 38.4853 ± 27.7240 | 50.6188 ± 30.0618 | 90.4429 ± 29.0305 | 127.2172 ± 74.1863 |
| MCP | 10.5857 ± 6.3202 | 17.3298 ± 11.1574 | 51.4537 ± 30.1220 | 43.3855 ± 33.6452 | - | - | - | - | - |
| MdLF_L | 14.3665 ± 8.3194 | 8.4157 ± 5.5276 | 56.8422 ± 49.5838 | 36.7739 ± 31.1822 | 0.4163 ± 0.1573 | 50.8289 ± 30.1481 | 61.8143 ± 23.6356 | 97.4048 ± 46.8142 | 125.1156 ± 66.5788 |
| MdLF_R | 10.7450 ± 7.9964 | 8.2246 ± 5.9636 | 55.3305 ± 50.2630 | 36.1239 ± 34.6153 | 0.4632 ± 0.1531 | 45.8801 ± 36.1392 | 71.4388 ± 28.5265 | 104.3784 ± 42.4640 | 135.6276 ± 68.1296 |
| OR_ML_L | 4.1400 ± 3.8627 | 10.0678 ± 5.7946 | 59.0603 ± 31.4590 | 42.5010 ± 24.2010 | 0.3573 ± 0.1703 | 45.0228 ± 31.4545 | 59.9386 ± 31.6911 | 95.1105 ± 36.5221 | 88.5938 ± 69.5327 |
| OR_ML_R | 5.5102 ± 7.1274 | 12.7676 ± 7.6223 | 86.4971 ± 42.1039 | 62.5457 ± 34.9648 | 0.2138 ± 0.1196 | 47.8760 ± 35.7274 | 53.2716 ± 34.1940 | 111.9585 ± 39.9669 | 69.9509 ± 58.2121 |
| POPT_L | 4.5857 ± 3.1448 | 5.1290 ± 3.8614 | 23.6069 ± 13.4042 | 21.2768 ± 11.9264 | 0.2908 ± 0.2180 | 144.0642 ± 64.5311 | 99.7508 ± 63.4215 | 121.5503 ± 77.3041 | 126.2808 ± 88.1962 |
| POPT_R | 4.4651 ± 3.0735 | 5.7789 ± 4.2919 | 26.3258 ± 19.7795 | 22.0600 ± 14.7985 | 0.3237 ± 0.1824 | 125.5377 ± 67.2363 | 113.6394 ± 65.2036 | 99.3050 ± 75.7996 | 153.7215 ± 75.8021 |
| PYT_L | 3.8525 ± 3.7411 | 6.0467 ± 3.6747 | 63.3981 ± 21.2531 | 36.9747 ± 14.4668 | 0.4169 ± 0.2487 | 165.0297 ± 48.0601 | 99.1033 ± 68.9259 | 145.0567 ± 67.5981 | 146.3560 ± 83.8815 |
| PYT_R | 3.6086 ± 3.4015 | 5.9533 ± 3.4104 | 63.3580 ± 17.7112 | 35.7914 ± 12.2064 | 0.3720 ± 0.2331 | 166.4927 ± 38.0673 | 79.4674 ± 65.5698 | 133.1660 ± 68.2547 | 155.3007 ± 75.5170 |
| SLF_L | 11.0423 ± 7.2426 | 9.7943 ± 7.3305 | 96.2303 ± 43.7705 | 66.0393 ± 36.0735 | 0.4064 ± 0.1902 | 79.7213 ± 77.7249 | 65.8042 ± 49.2349 | 152.9229 ± 33.4092 | 121.1710 ± 75.6789 |
| SLF_R | 6.9994 ± 5.5779 | 11.6982 ± 6.1827 | 68.6048 ± 47.6555 | 47.0659 ± 33.1078 | 0.5068 ± 0.1499 | 53.3537 ± 31.5136 | 74.2599 ± 49.0805 | 113.8505 ± 43.8950 | 126.6906 ± 70.0115 |
| UF_L | 12.1606 ± 8.3958 | 14.4482 ± 11.9936 | 77.7074 ± 49.4321 | 43.2770 ± 33.8591 | 0.4197 ± 0.1874 | 36.2327 ± 46.3920 | 62.0342 ± 59.2523 | 61.0680 ± 57.6517 | 108.2708 ± 90.1743 |
| UF_R | 5.9925 ± 4.5454 | 13.8315 ± 10.2998 | 59.7495 ± 47.9775 | 36.6240 ± 34.8001 | 0.3359 ± 0.2197 | 69.6248 ± 57.9120 | 76.1830 ± 57.7207 | 62.0682 ± 57.6067 | 122.7808 ± 84.0423 |

**Tab. S11 | Geometry and connectivity of bundles generated with TractSeg.** Absolute percent differences are presented as mean ± standard deviation. A dash '−' indicates an absent bundle.

| Bundles | Geometry APD (↓) | | | | Pearson (↑) | Connectivity APD (↓) | | | |
|---|---|---|---|---|---|---|---|---|---|
| | Mean Length | Span | Volume | Surface Area | | Density | CPL | Efficiency | Modularity |
| AF_L | 25.9307 ± 7.1867 | 24.2242 ± 16.2981 | 31.8824 ± 21.0098 | 20.4990 ± 16.9734 | 0.3509 ± 0.1771 | 130.7930 ± 72.4398 | 159.0896 ± 43.0805 | 164.1861 ± 44.4708 | 150.2655 ± 65.7205 |
| AF_R | 14.7723 ± 7.3218 | 11.0559 ± 7.5228 | 52.7674 ± 32.2260 | 40.3864 ± 28.2097 | 0.3988 ± 0.1670 | 125.1083 ± 79.8707 | 145.3272 ± 56.5834 | 167.6320 ± 36.8620 | 160.5039 ± 60.8569 |
| CG_L | 18.7440 ± 12.7914 | 14.2218 ± 10.0264 | 38.7323 ± 28.7776 | 47.8650 ± 23.4019 | 0.3249 ± 0.1296 | 135.2870 ± 75.4661 | 141.3610 ± 70.0386 | 170.3936 ± 37.8569 | 152.3122 ± 82.8314 |
| CG_R | 22.5000 ± 10.5192 | 16.5651 ± 10.0830 | 33.9811 ± 27.1874 | 36.3449 ± 23.2912 | 0.3087 ± 0.1261 | 141.5595 ± 66.1656 | 142.2118 ± 58.8784 | 180.0850 ± 23.3998 | 136.0995 ± 80.5869 |
| FPT_L | 13.4802 ± 7.2197 | 9.5510 ± 8.4750 | 64.5067 ± 30.8731 | 57.5199 ± 26.4426 | 0.1680 ± 0.2294 | 158.9462 ± 63.2939 | 141.9863 ± 58.4294 | 160.8648 ± 62.5728 | 114.9763 ± 97.4752 |
| FPT_R | 11.5853 ± 7.1216 | 8.0825 ± 8.3687 | 67.1759 ± 27.5980 | 63.1457 ± 22.2177 | 0.2598 ± 0.1107 | 152.9840 ± 75.4014 | 133.0723 ± 54.7477 | 166.1270 ± 57.0859 | 150.7430 ± 84.0820 |
| IFOF_L | 26.7511 ± 5.9866 | 22.5532 ± 7.7486 | 86.7965 ± 22.1776 | 66.7327 ± 26.5753 | 0.5757 ± 0.1183 | 177.6029 ± 48.6158 | 176.1019 ± 45.8698 | 191.9643 ± 19.3959 | 153.1065 ± 76.6459 |
| IFOF_R | 20.6587 ± 9.8153 | 15.1711 ± 8.8905 | 86.3342 ± 27.3037 | 59.7589 ± 24.1881 | 0.6215 ± 0.0737 | 172.7907 ± 54.3458 | 190.6885 ± 7.7336 | 192.0111 ± 15.6011 | 181.3627 ± 51.0550 |
| ILF_L | 13.7164 ± 7.1017 | 9.9979 ± 5.2927 | 104.9650 ± 32.9202 | 97.1996 ± 28.6238 | 0.2580 ± 0.1279 | 143.1780 ± 74.5215 | 124.8479 ± 65.6551 | 146.1193 ± 68.6983 | 112.3669 ± 89.6710 |
| ILF_R | 15.4987 ± 5.2703 | 11.1992 ± 4.9452 | 105.8440 ± 23.1539 | 100.1605 ± 18.1797 | 0.2536 ± 0.1253 | 148.1518 ± 66.3389 | 127.4731 ± 66.1078 | 137.1687 ± 76.4524 | 116.5496 ± 88.0632 |
| MCP | 23.5557 ± 10.4028 | 18.2712 ± 10.1143 | 27.4852 ± 20.7930 | 31.6378 ± 20.6518 | - | - | - | - | - |
| MdLF_L | 24.7503 ± 11.2406 | 27.8340 ± 11.2193 | 59.1514 ± 53.3226 | 76.3347 ± 37.5720 | 0.1647 ± 0.1745 | 140.1730 ± 60.7228 | 153.2225 ± 45.6543 | 169.1587 ± 32.9949 | 166.2077 ± 50.8270 |
| MdLF_R | 22.8927 ± 10.1179 | 23.0922 ± 12.9674 | 54.8791 ± 53.7793 | 70.3347 ± 38.4399 | 0.2992 ± 0.1727 | 133.8281 ± 72.7731 | 140.3029 ± 56.5685 | 167.3518 ± 37.8551 | 150.1924 ± 71.9429 |
| POPT_L | 14.8860 ± 7.2696 | 14.0702 ± 8.4787 | 50.0688 ± 26.9983 | 36.6492 ± 19.9329 | 0.1577 ± 0.1431 | 136.1473 ± 77.0702 | 163.3863 ± 46.7693 | 158.3369 ± 58.5575 | 138.0840 ± 77.3743 |
| POPT_R | 17.1744 ± 5.8879 | 16.5471 ± 7.4166 | 46.3896 ± 26.6055 | 35.9621 ± 22.3454 | 0.0783 ± 0.0640 | 136.9786 ± 79.0322 | 180.9065 ± 15.7748 | 157.1115 ± 60.3991 | 141.0503 ± 67.6567 |
| UF_L | 51.4600 ± 13.1133 | 53.1904 ± 17.0309 | 60.3029 ± 34.1972 | 78.7776 ± 30.5903 | 0.2849 ± 0.1600 | 110.6975 ± 91.1430 | 121.1829 ± 77.2268 | 114.8551 ± 88.6648 | 74.8395 ± 92.6888 |
| UF_R | 47.0855 ± 16.1410 | 49.4637 ± 18.4856 | 60.7744 ± 30.4797 | 80.1602 ± 30.7219 | 0.3391 ± 0.2352 | 115.4410 ± 87.1842 | 123.9890 ± 75.5244 | 132.9097 ± 73.7247 | 114.4904 ± 89.8244 |

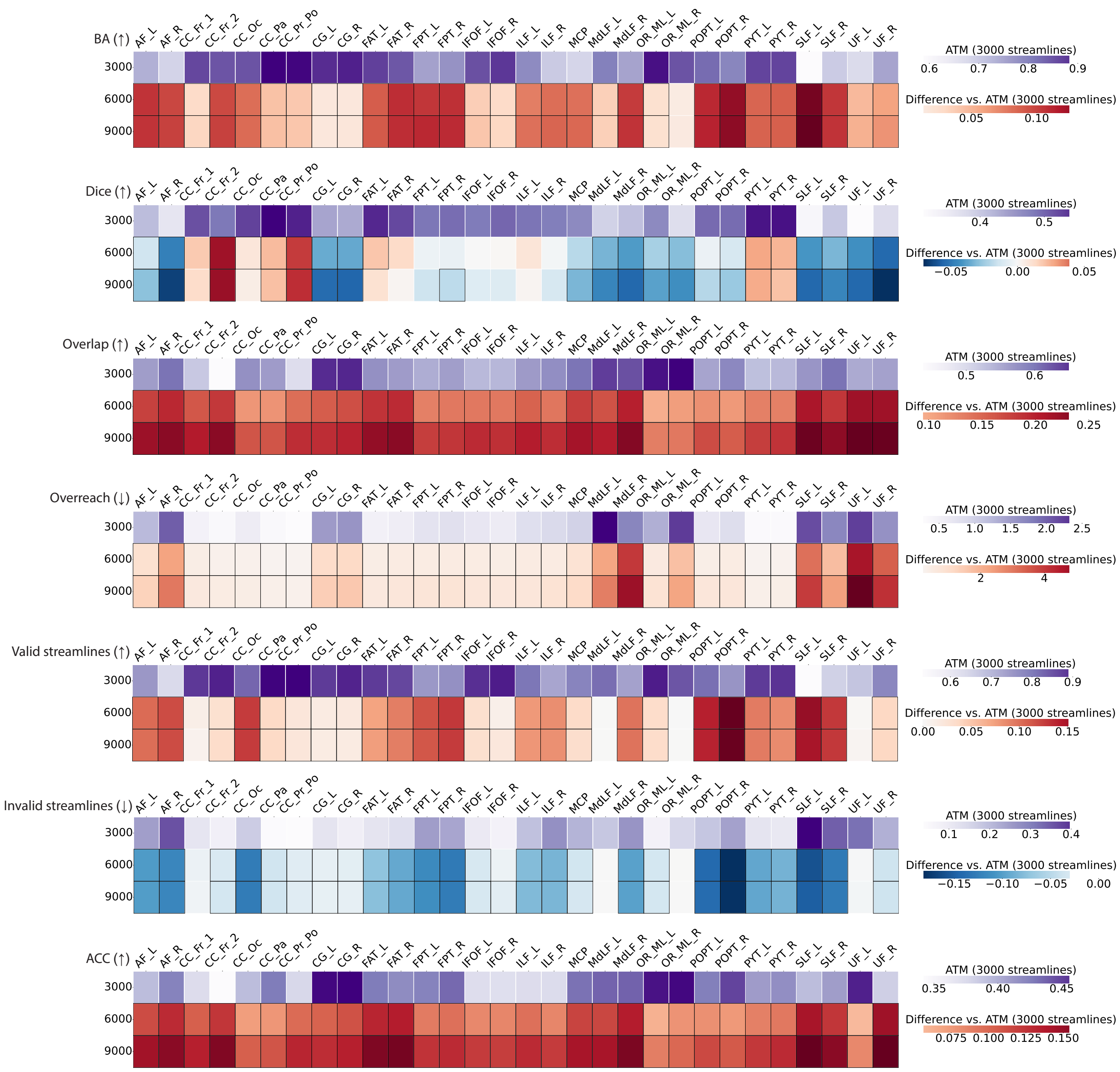

**Fig. S12 | Similarity and coverage of bundles generated with ATM using 3000, 6000, and 9000 streamlines per bundle.** The first row of each heatmap shows the values for 3,000 streamlines per bundle as the baseline. The subsequent rows display the paired performance differences of ATM with other streamline counts relative to this baseline. The arrow indicates preference: ↑ higher is better, ↓ lower is better. Cells outlined with squares indicate statistically significant differences.

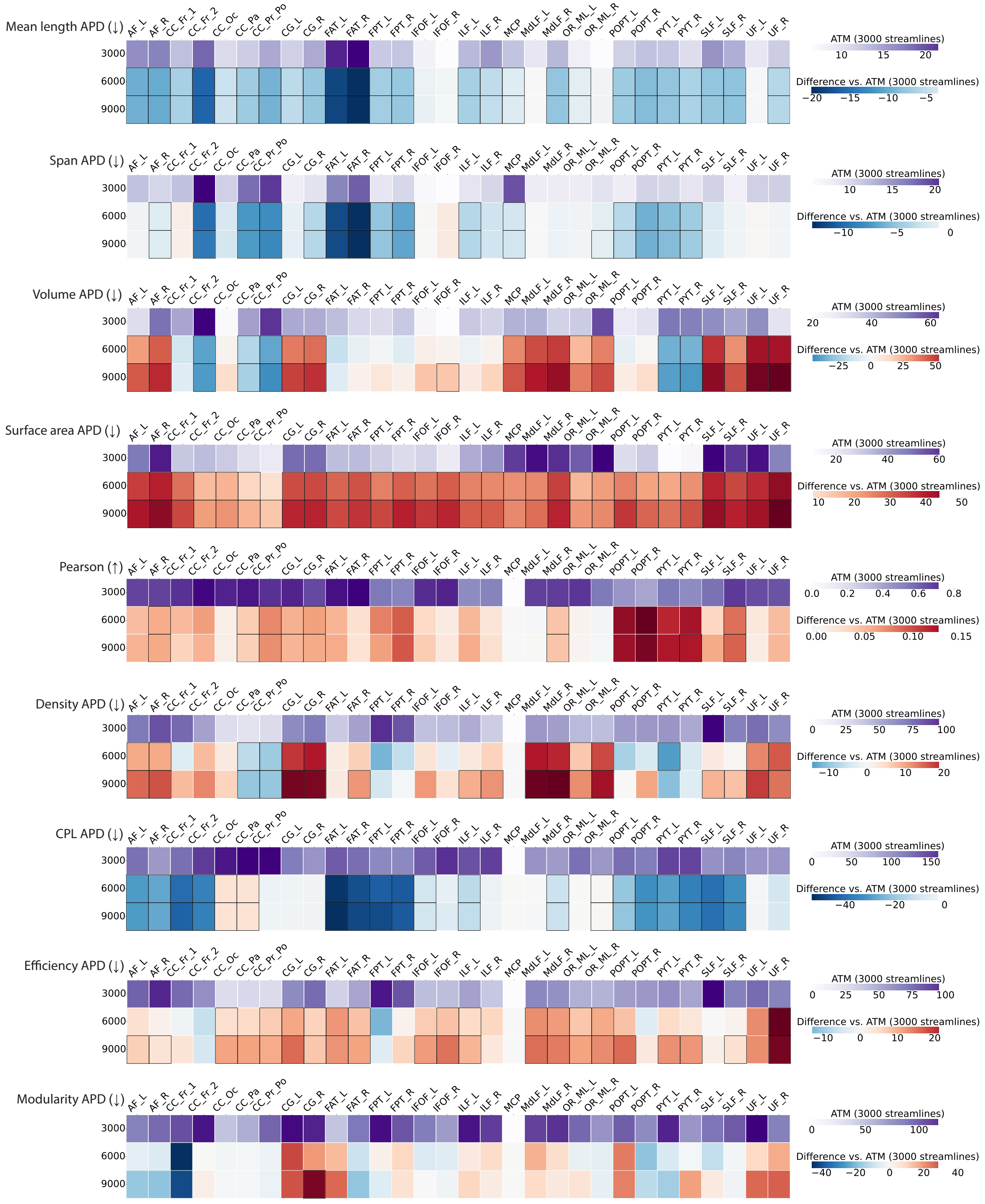


**Fig. S13 | Geometry and connectivity of bundles generated with ATM using 3,000, 6,000, and 9,000 streamlines per bundle.** The first row of each heatmap shows the values for 3,000 streamlines per bundle as the baseline. The subsequent rows display the paired performance differences of ATM with other streamline counts relative to this baseline. The arrow indicates preference: ↑ higher is better, ↓ lower is better. Cells outlined with squares indicate statistically significant differences.

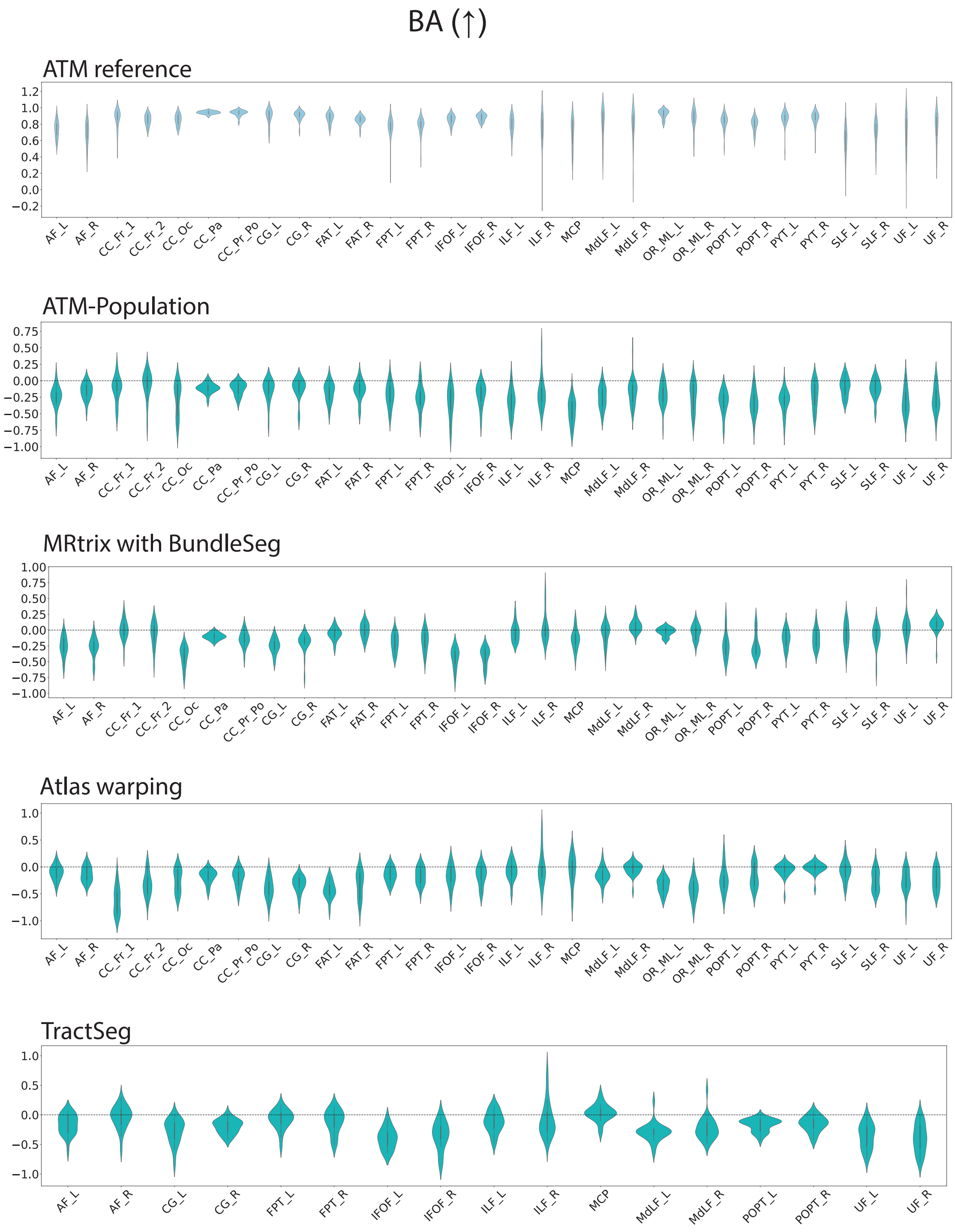


**Fig. S14 | BA of bundles generated with ATM, ATM-Population, MRtrix with BundleSeg, SCIL WM atlas warping, and TractSeg.** The first violin plot shows the ATM values as the baseline. The subsequent rows display the paired performance differences of other methods relative to this baseline. For each metric, the arrow indicates preference: ↑ higher is better and ↓ lower is better.

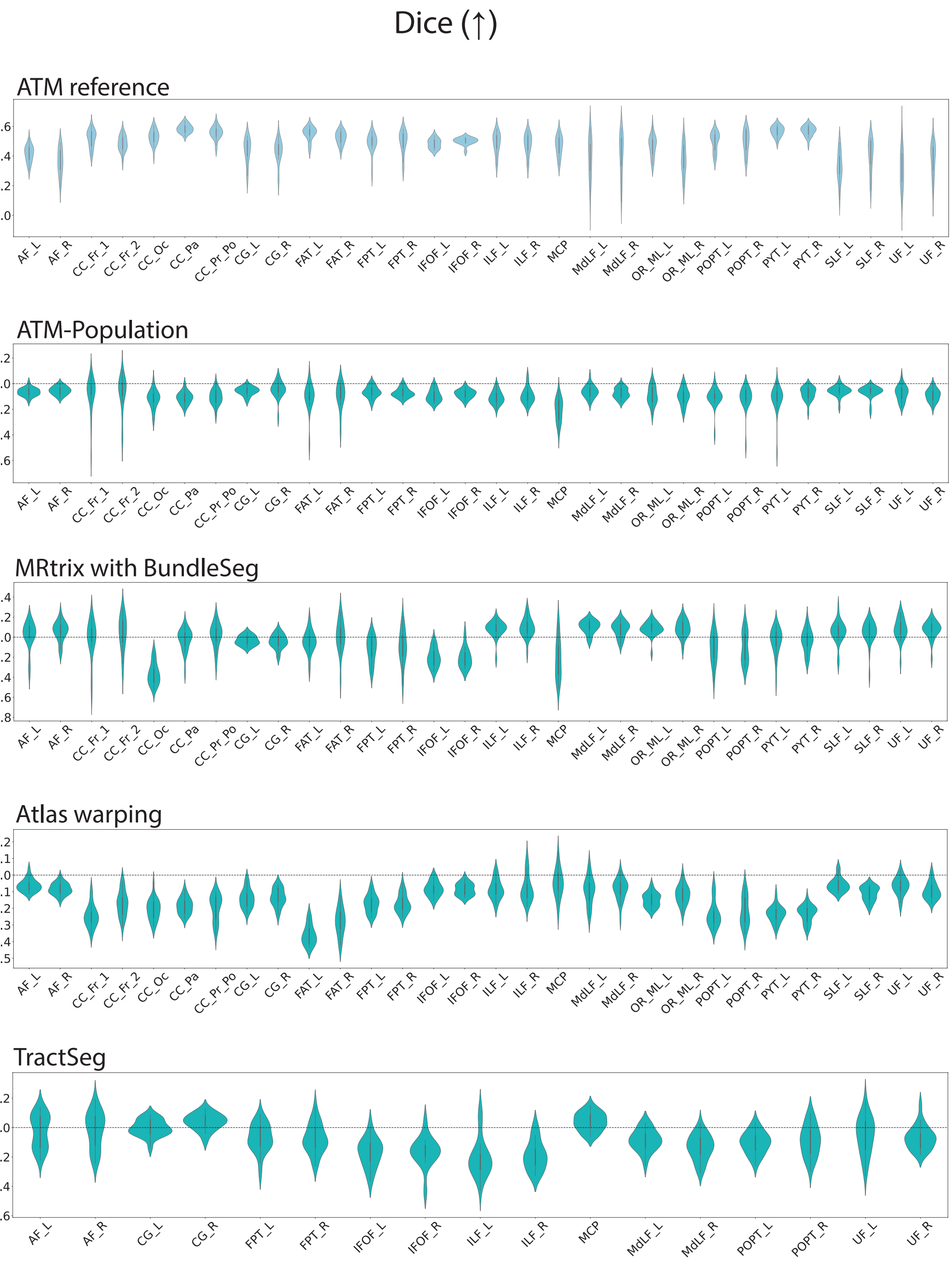


**Fig. S15 | Dice scores of bundles generated with ATM, ATM-Population, MRtrix with BundleSeg, SCIL WM atlas warping, and TractSeg.** The first violin plot shows the ATM values as the baseline. The subsequent rows display the paired performance differences of other methods relative to this baseline. For each metric, the arrow indicates preference: ↑ higher is better and ↓ lower is better.

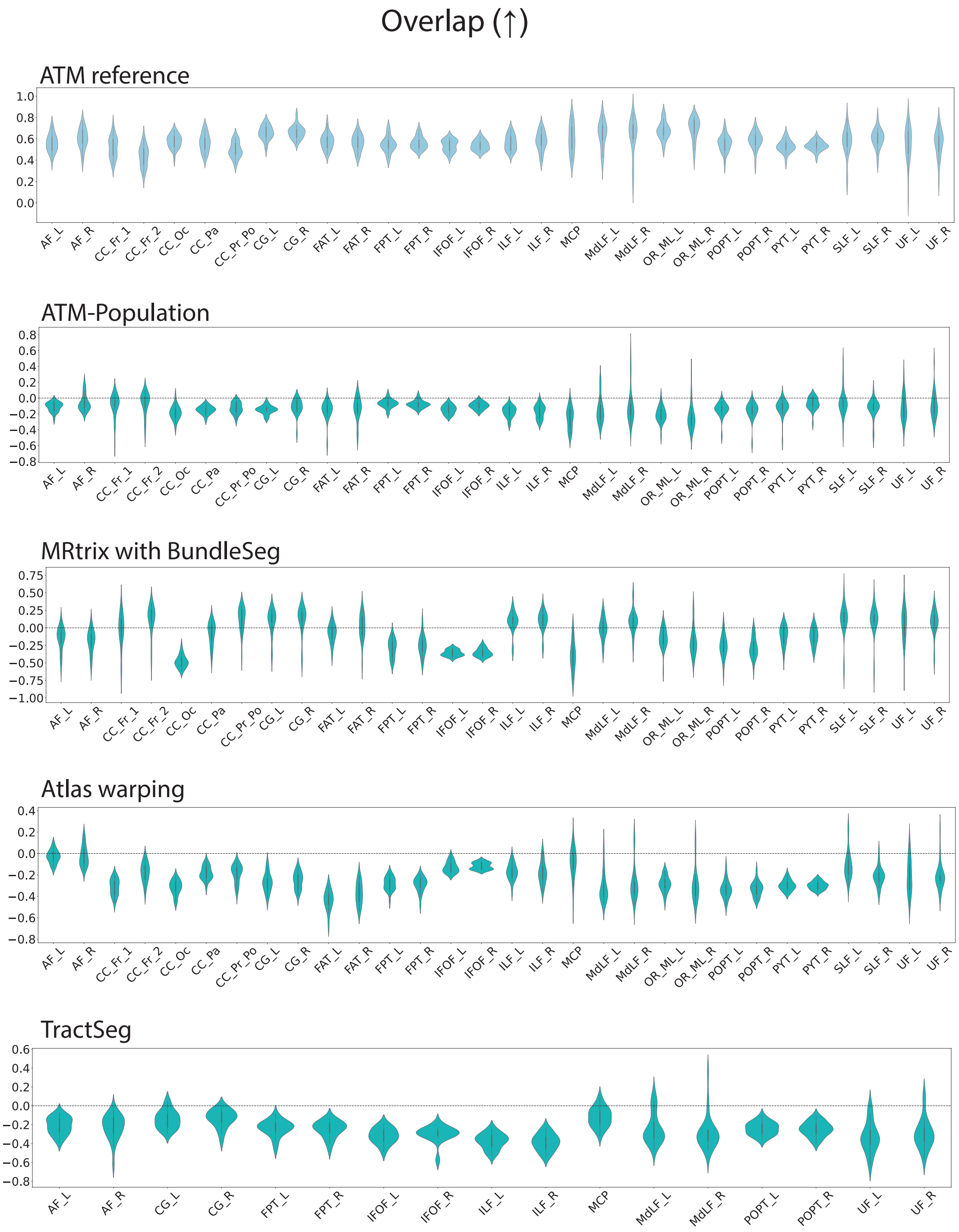


**Fig. S16 | Overlap of bundles generated with ATM, ATM-Population, MRtrix, SCIL WM atlas warping, and TractSeg.** The first violin plot shows the ATM values as the baseline. The subsequent rows display the paired performance differences of other methods relative to this baseline. For each metric, the arrow indicates preference: ↑ higher is better and ↓ lower is better.

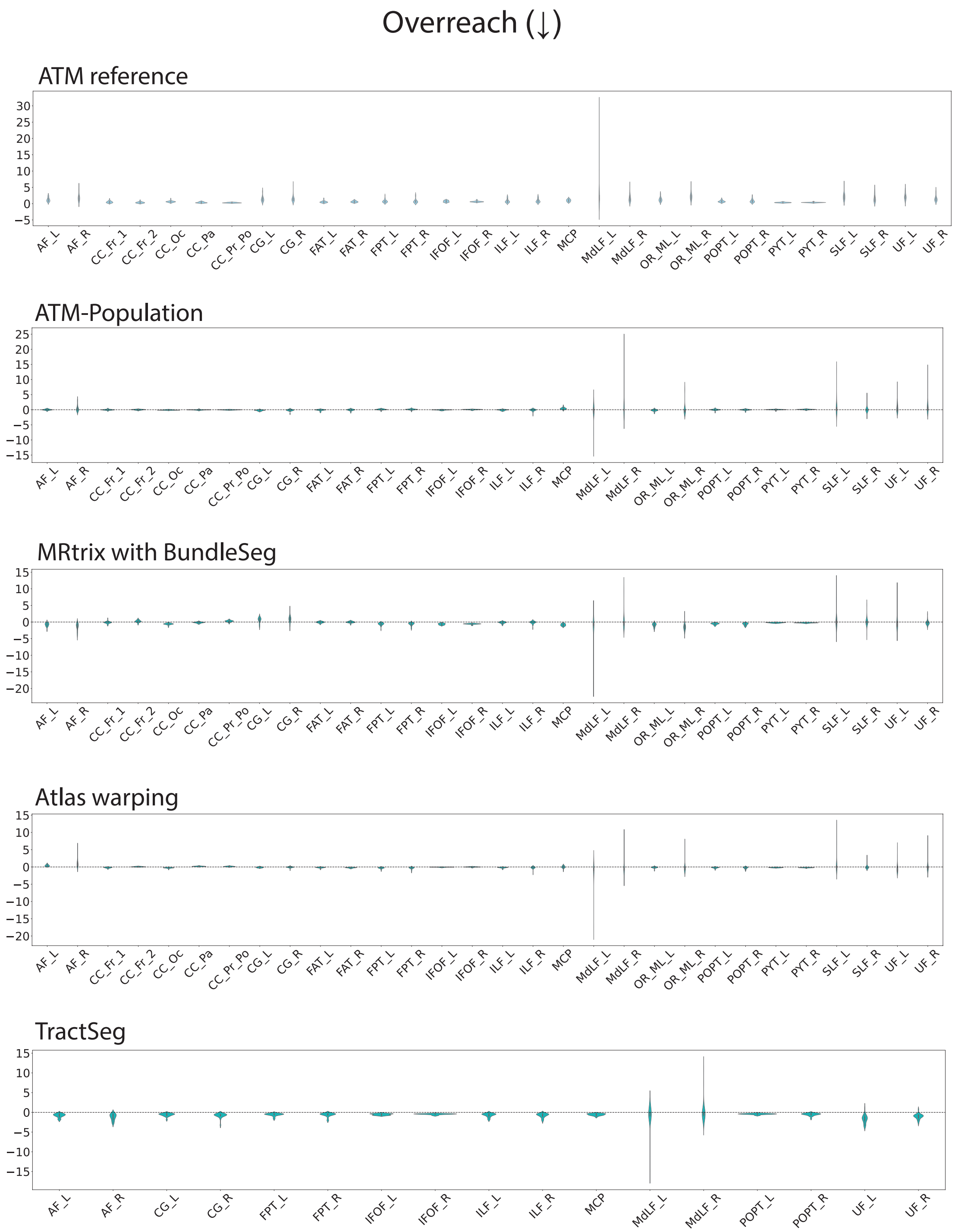


**Fig. S17 | Overreach of bundles generated with ATM, ATM-Population, MRtrix with BundleSeg, SCIL WM atlas warping, and TractSeg.** The first violin plot shows the ATM values as the baseline. The subsequent rows display the paired performance differences of other methods relative to this baseline. For each metric, the arrow indicates preference: ↑ higher is better and ↓ lower is better.

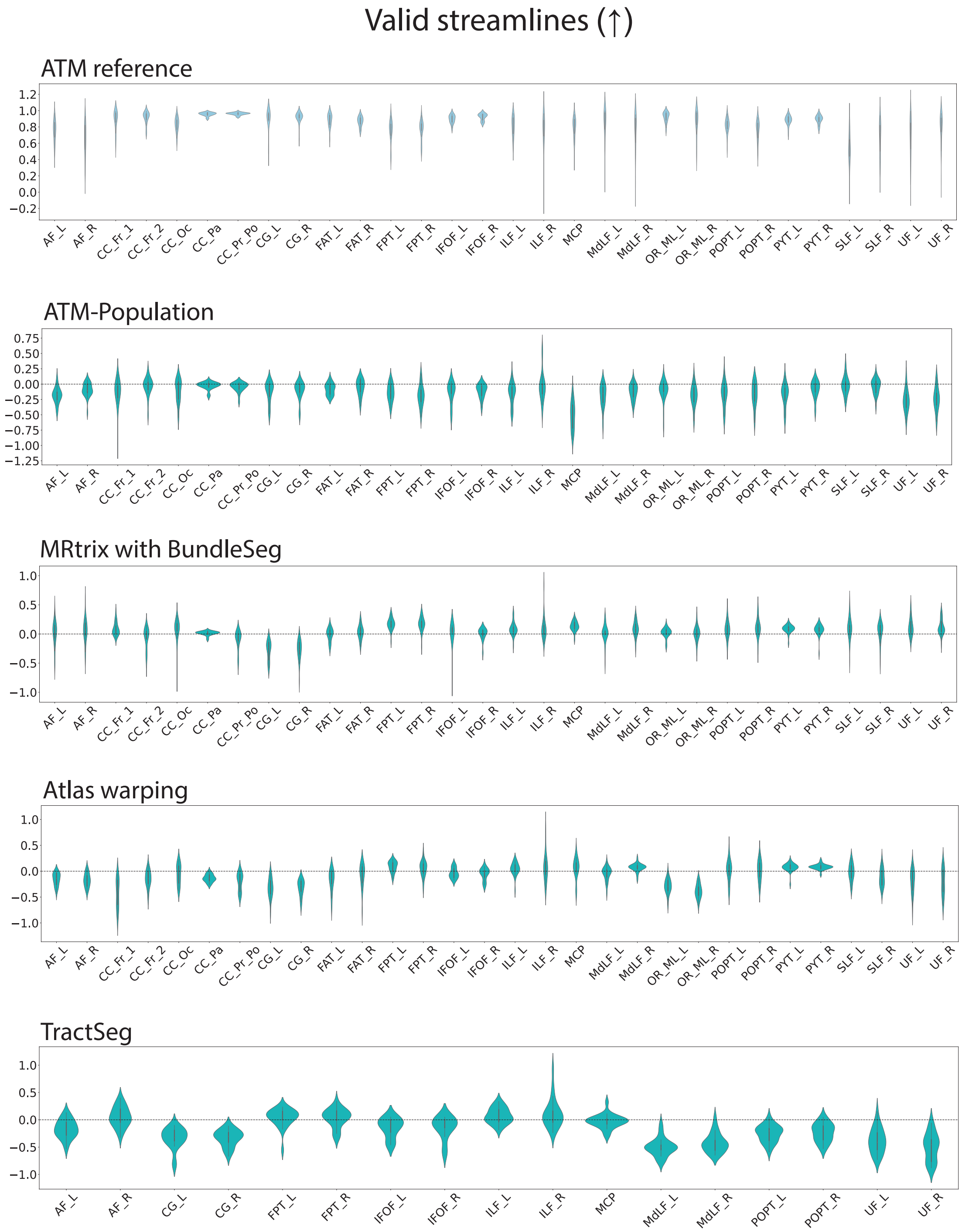


**Fig. S18 | Ratio of valid streamlines in bundles generated with ATM, ATM-Population, MRtrix with BundleSeg, SCIL WM atlas warping, and TractSeg.** The first violin plot shows the ATM values as the baseline. The subsequent rows display the paired performance differences of other methods relative to this baseline. For each metric, the arrow indicates preference: ↑ higher is better and ↓ lower is better.

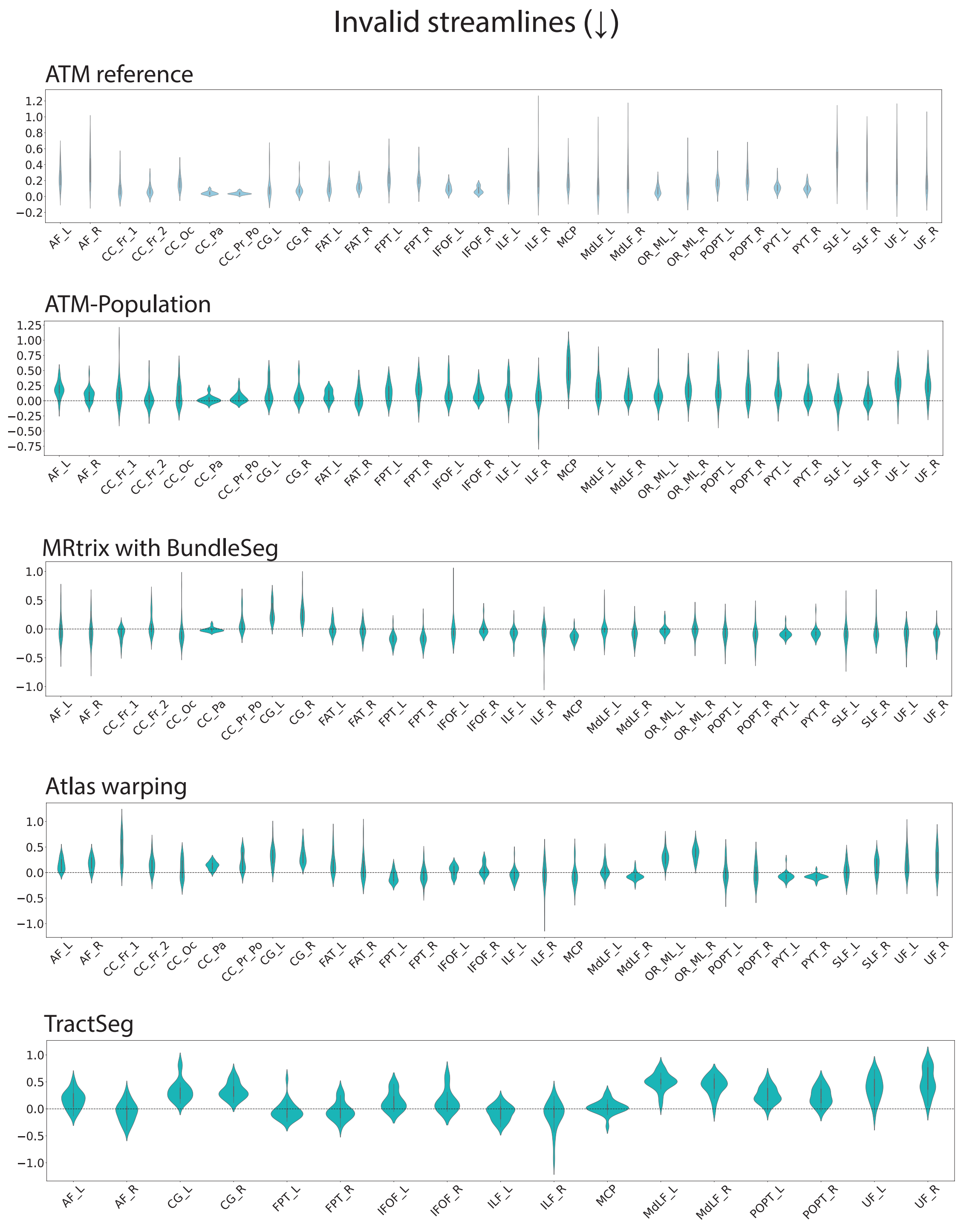


**Fig. S19 | Ratio of invalid streamlines in bundles generated with ATM, ATM-Population, MRtrix with BundleSeg, SCIL WM atlas warping, and TractSeg.** The first violin plot shows the ATM values as the baseline. The subsequent rows display the paired performance differences of other methods relative to this baseline. For each metric, the arrow indicates preference: ↑ higher is better and ↓ lower is better.

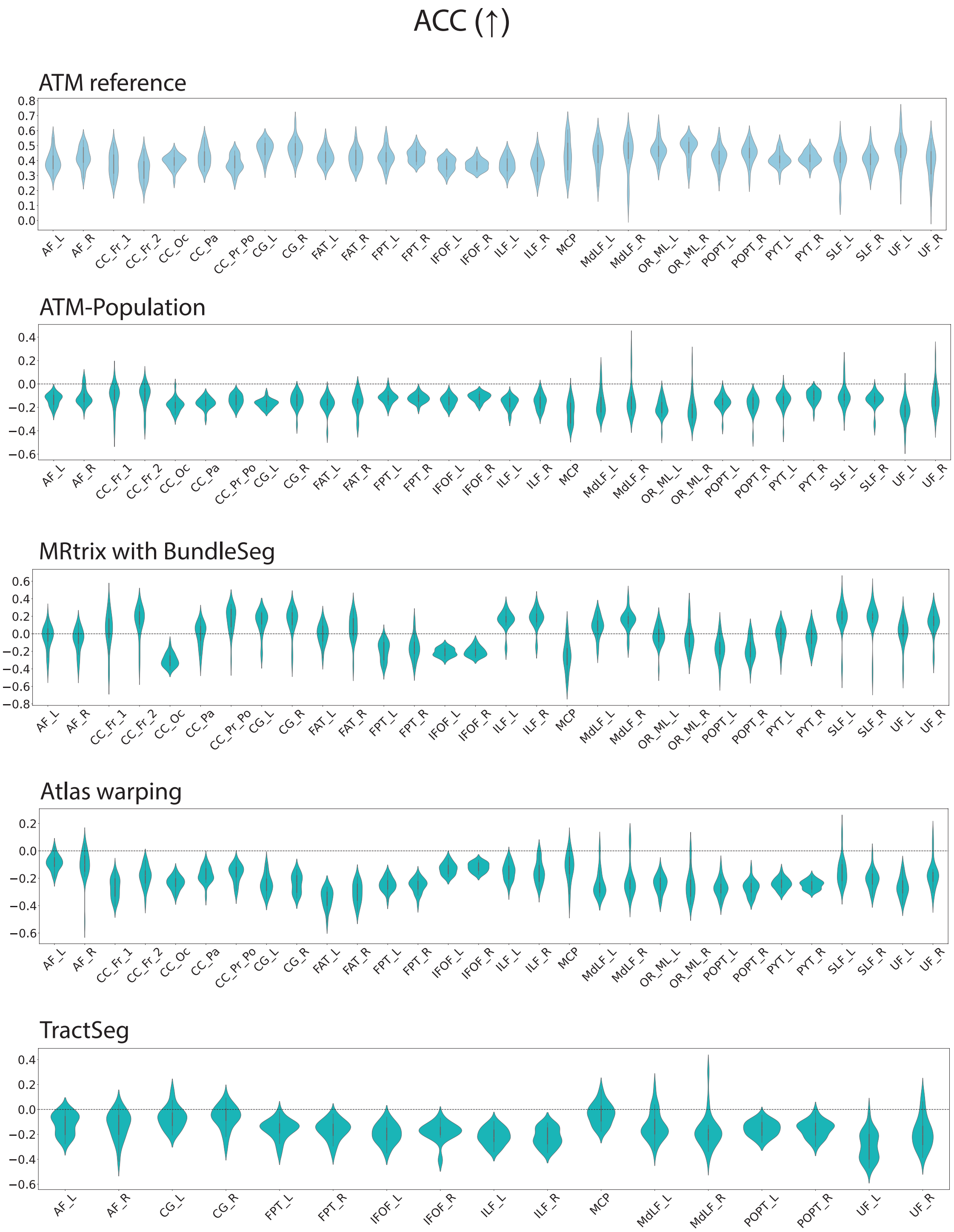


**Fig. S20 | Angular correlation coefficient of bundles generated with ATM, ATM-Population, MRtrix with BundleSeg, SCIL WM atlas warping, and TractSeg.** The first violin plot shows the ATM values as the baseline. The subsequent rows display the paired performance differences of other methods relative to this baseline. For each metric, the arrow indicates preference: ↑ higher is better and ↓ lower is better.

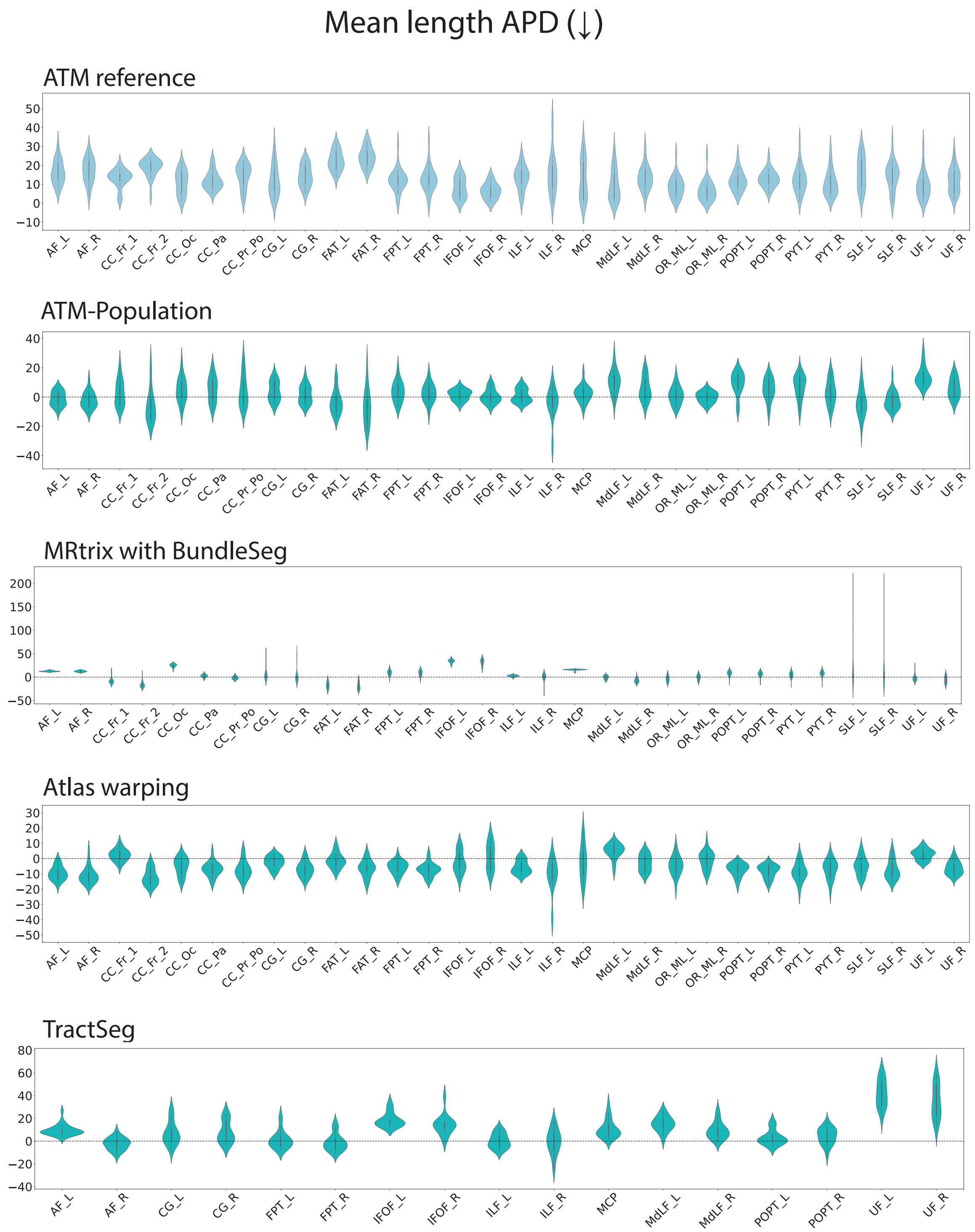


**Fig. S21 | Mean length APD of bundles generated with ATM, ATM-Population, MRtrix with BundleSeg, SCIL WM atlas warping, and TractSeg.** The first violin plot shows the ATM values as the baseline. The subsequent rows display the paired performance differences of other methods relative to this baseline. For each metric, the arrow indicates preference: ↑ higher is better and ↓ lower is better.

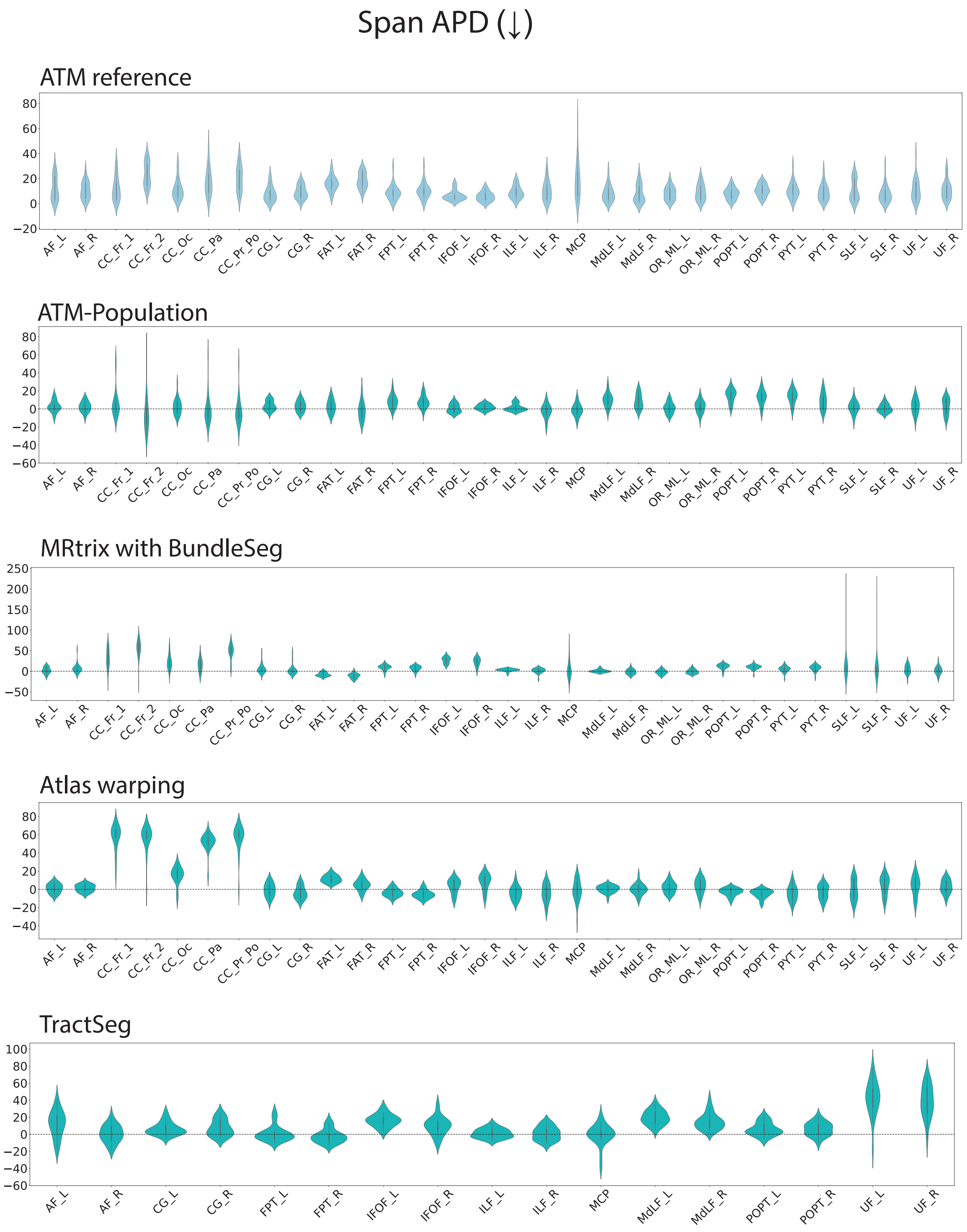


**Fig. S22 | Span APD of bundles generated with ATM, ATM-Population, MRtrix with BundleSeg, SCIL WM atlas warping, and TractSeg.** The first violin plot shows the ATM values as the baseline. The subsequent rows display the paired performance differences of other methods relative to this baseline. For each metric, the arrow indicates preference: ↑ higher is better and ↓ lower is better.

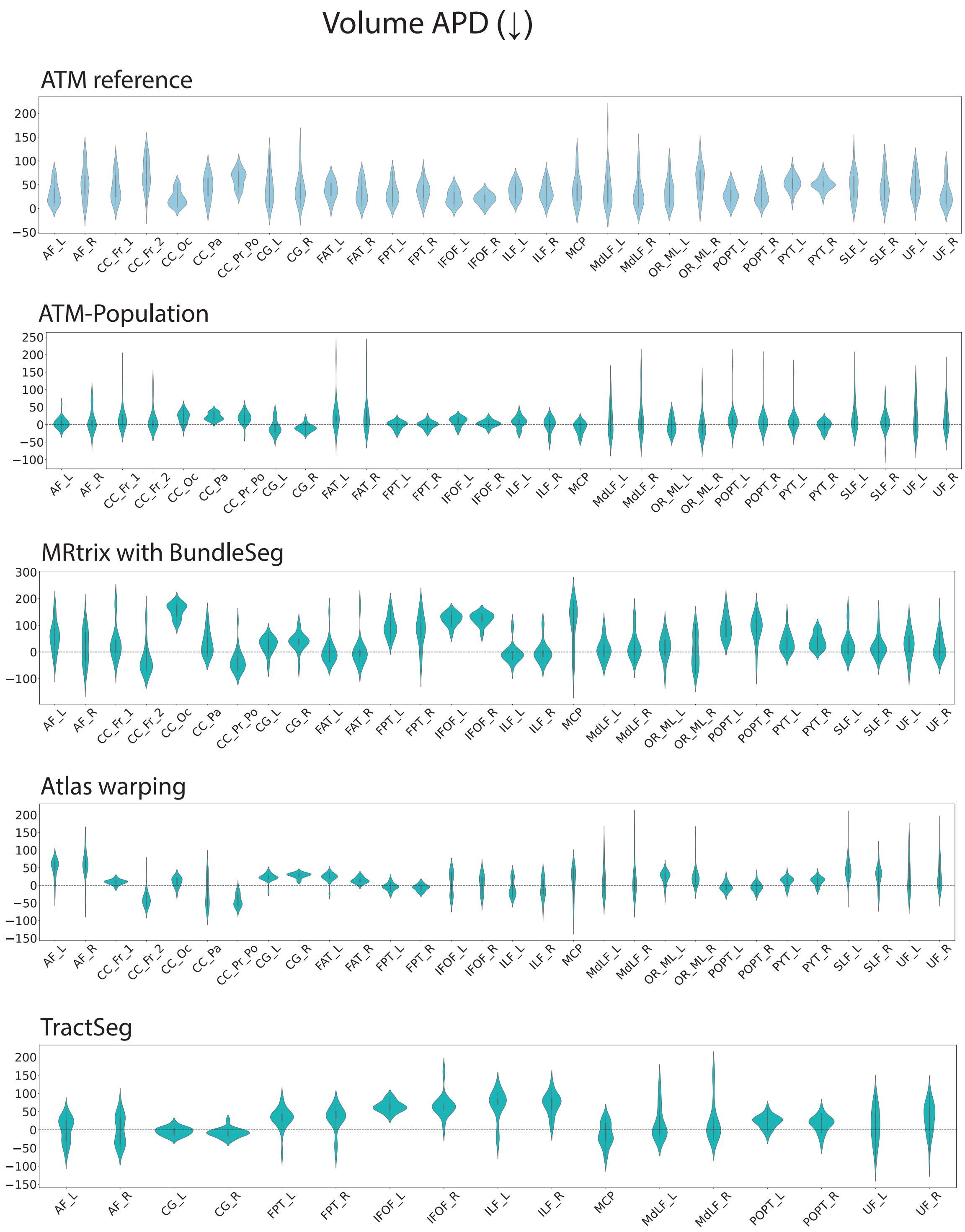


**Fig. S23 | Volume APD of bundles generated with ATM, ATM-Population, MRtrix with BundleSeg, SCIL WM atlas warping, and TractSeg.** The first violin plot shows the ATM values as the baseline. The subsequent rows display the paired performance differences of other methods relative to this baseline. For each metric, the arrow indicates preference: ↑ higher is better and ↓ lower is better.

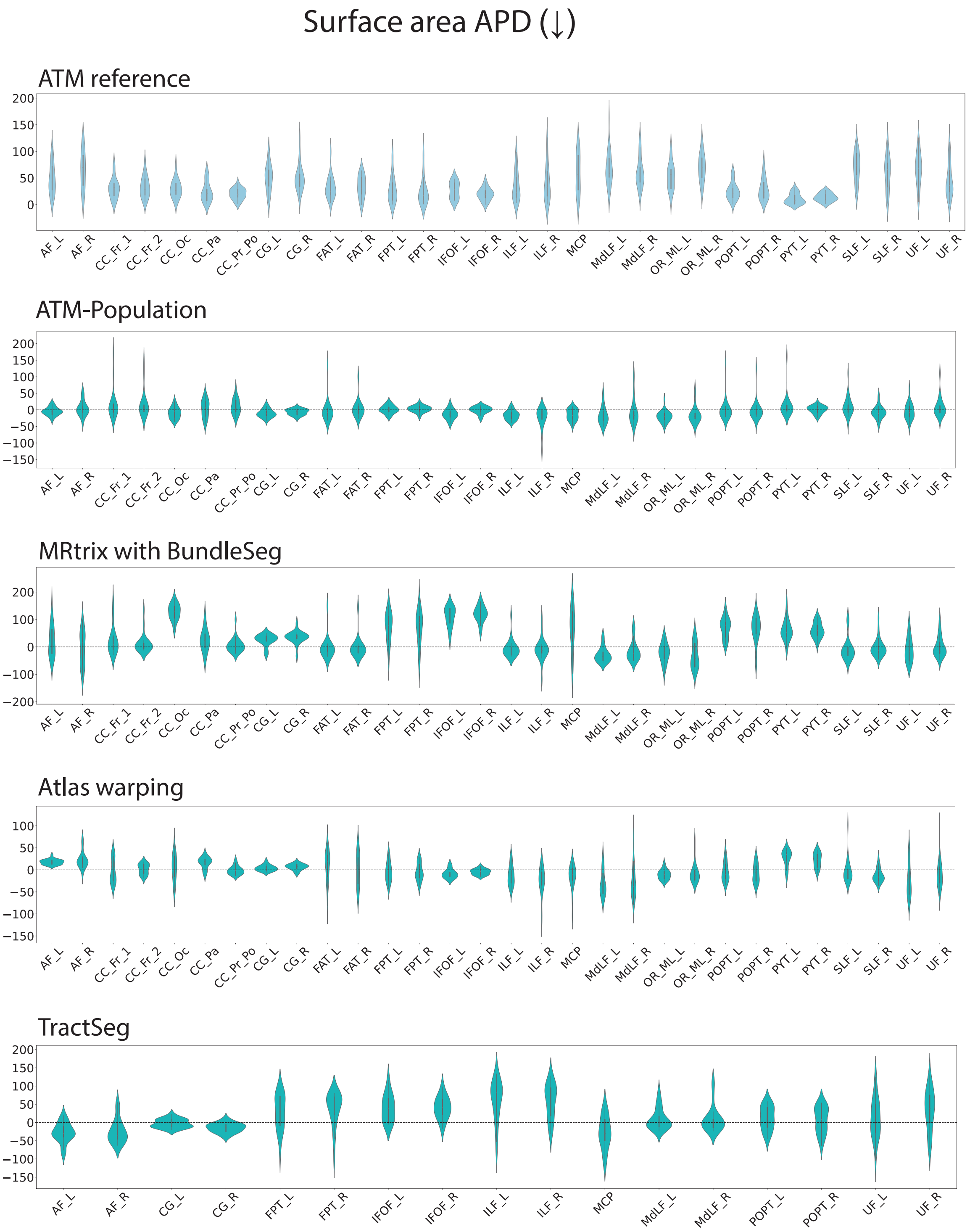


**Fig. S24 | Surface area APD of bundles generated with ATM, ATM-Population, MRtrix with BundleSeg, SCIL WM atlas warping, and TractSeg.** The first violin plot shows the ATM values as the baseline. The subsequent rows display the paired performance differences of other methods relative to this baseline. For each metric, the arrow indicates preference: ↑ higher is better and ↓ lower is better.

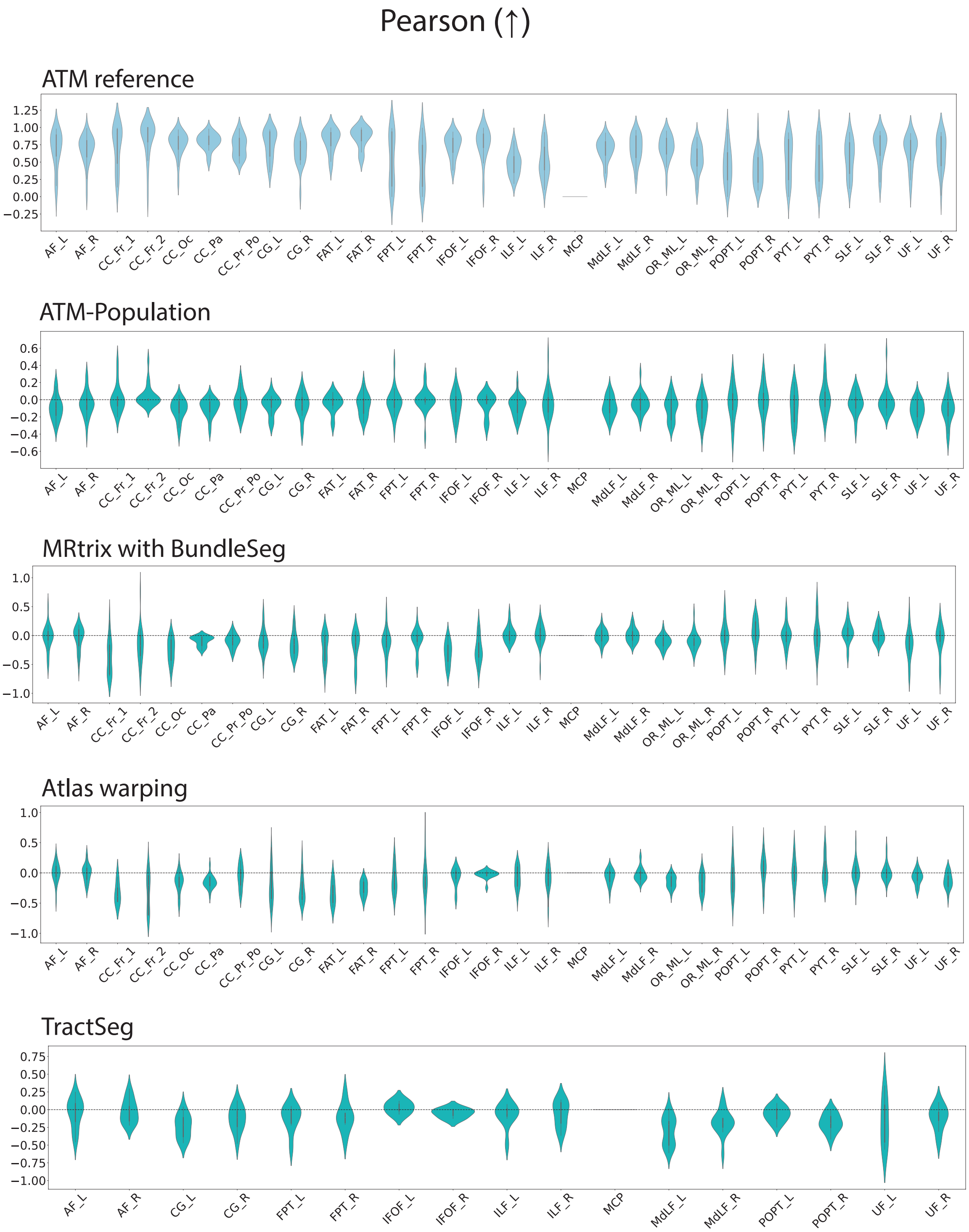


**Fig. S25 | Pearson corerlation of bundle connectomes generated with ATM, ATM-Population, MRtrix with BundleSeg, SCIL WM atlas warping, and TractSeg.** The first violin plot shows the ATM values as the baseline. The subsequent rows display the paired performance differences of other methods relative to this baseline. For each metric, the arrow indicates preference: ↑ higher is better and ↓ lower is better.

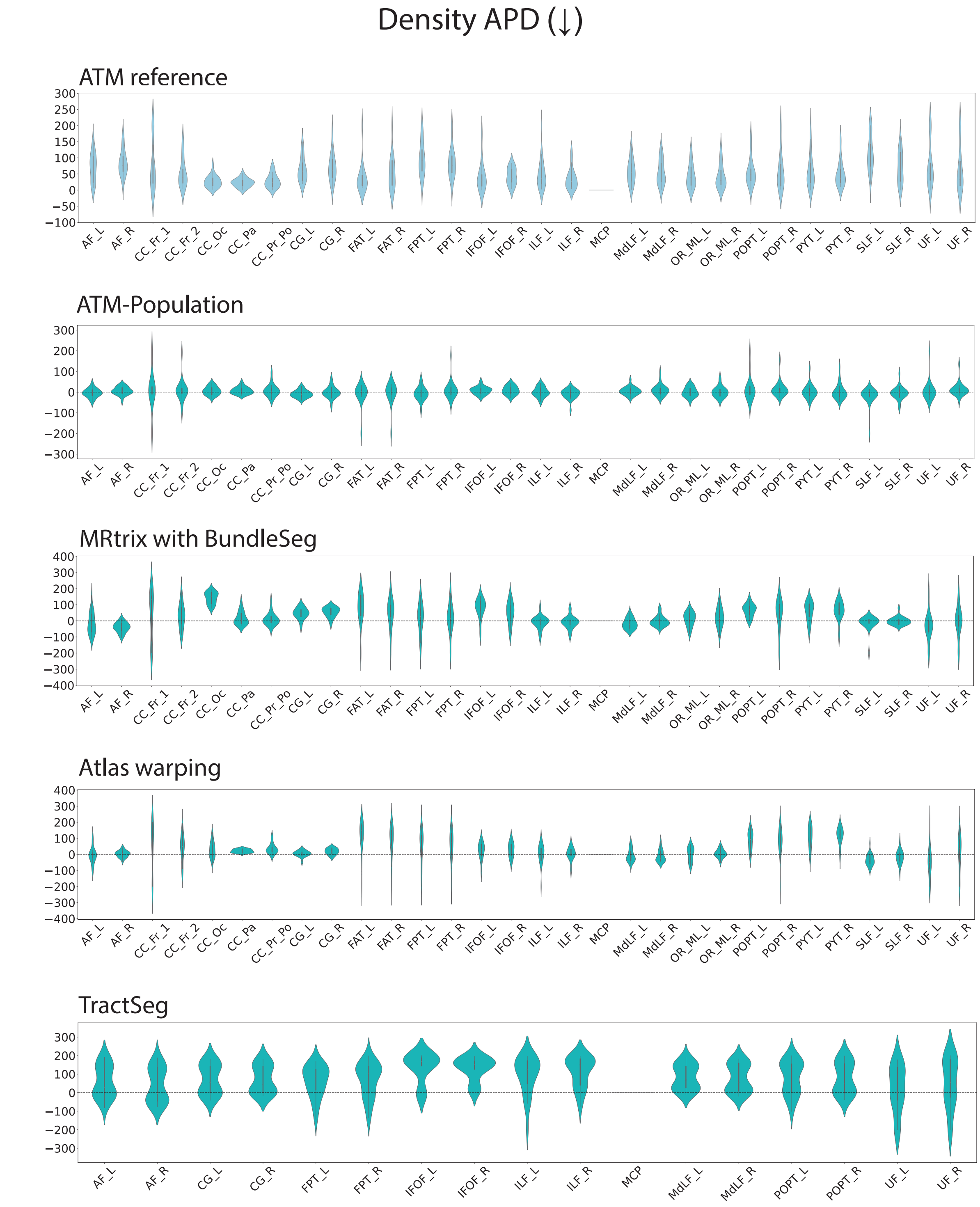


**Fig. S26 | Network density APD of bundle connectomes generated with ATM, ATM-Population, MRtrix with BundleSeg, SCIL WM atlas warping, and TractSeg.** The first violin plot shows the ATM values as the baseline. The subsequent rows display the paired performance differences of other methods relative to this baseline. For each metric, the arrow indicates preference: ↑ higher is better and ↓ lower is better.

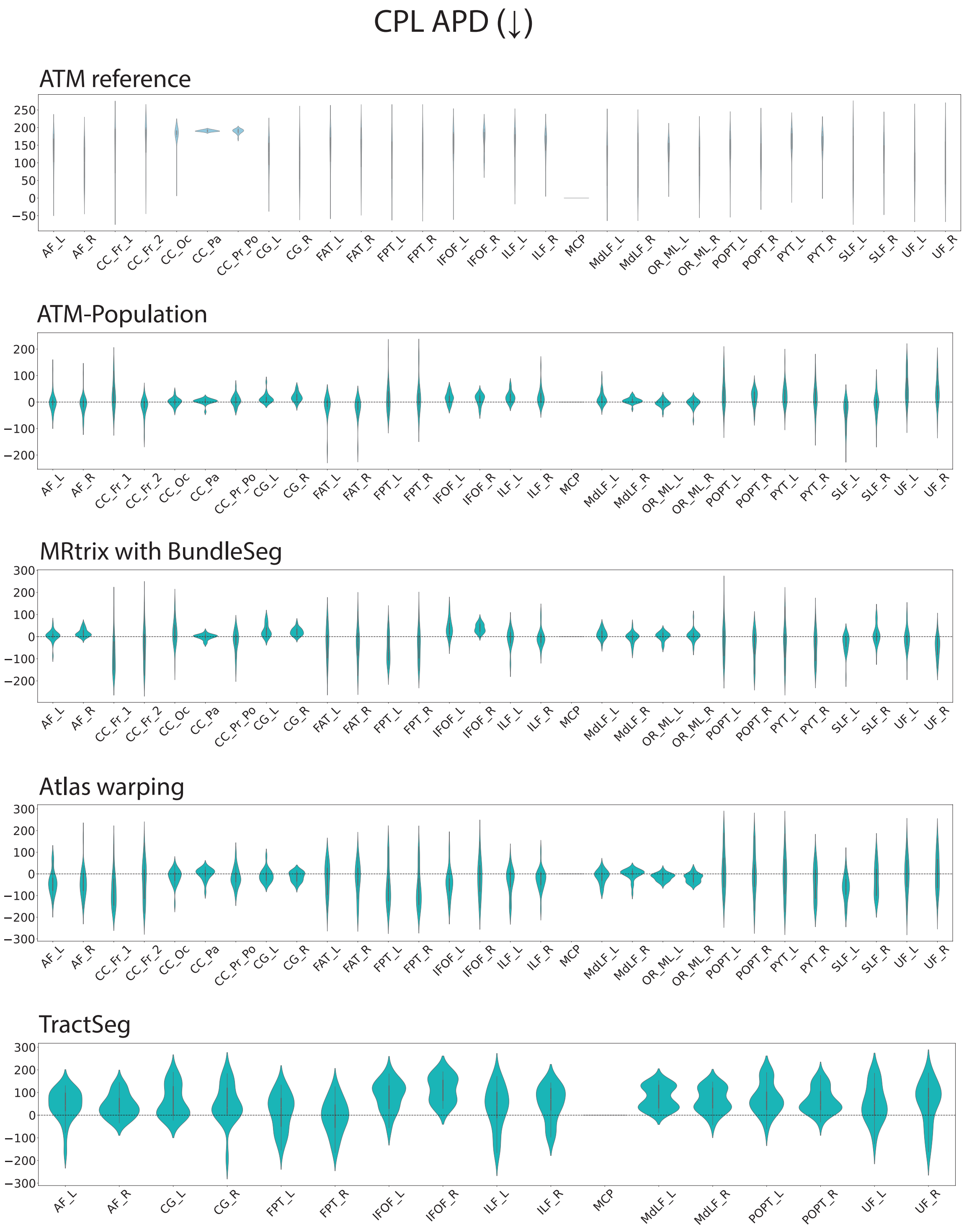


**Fig. S27 | Network CPL APD of bundle connectomes generated with ATM, ATM-Population, MRtrix with BundleSeg, SCIL WM atlas warping, and TractSeg.** The first violin plot shows the ATM values as the baseline. The subsequent rows display the paired performance differences of other methods relative to this baseline. For each metric, the arrow indicates preference: ↑ higher is better and ↓ lower is better.

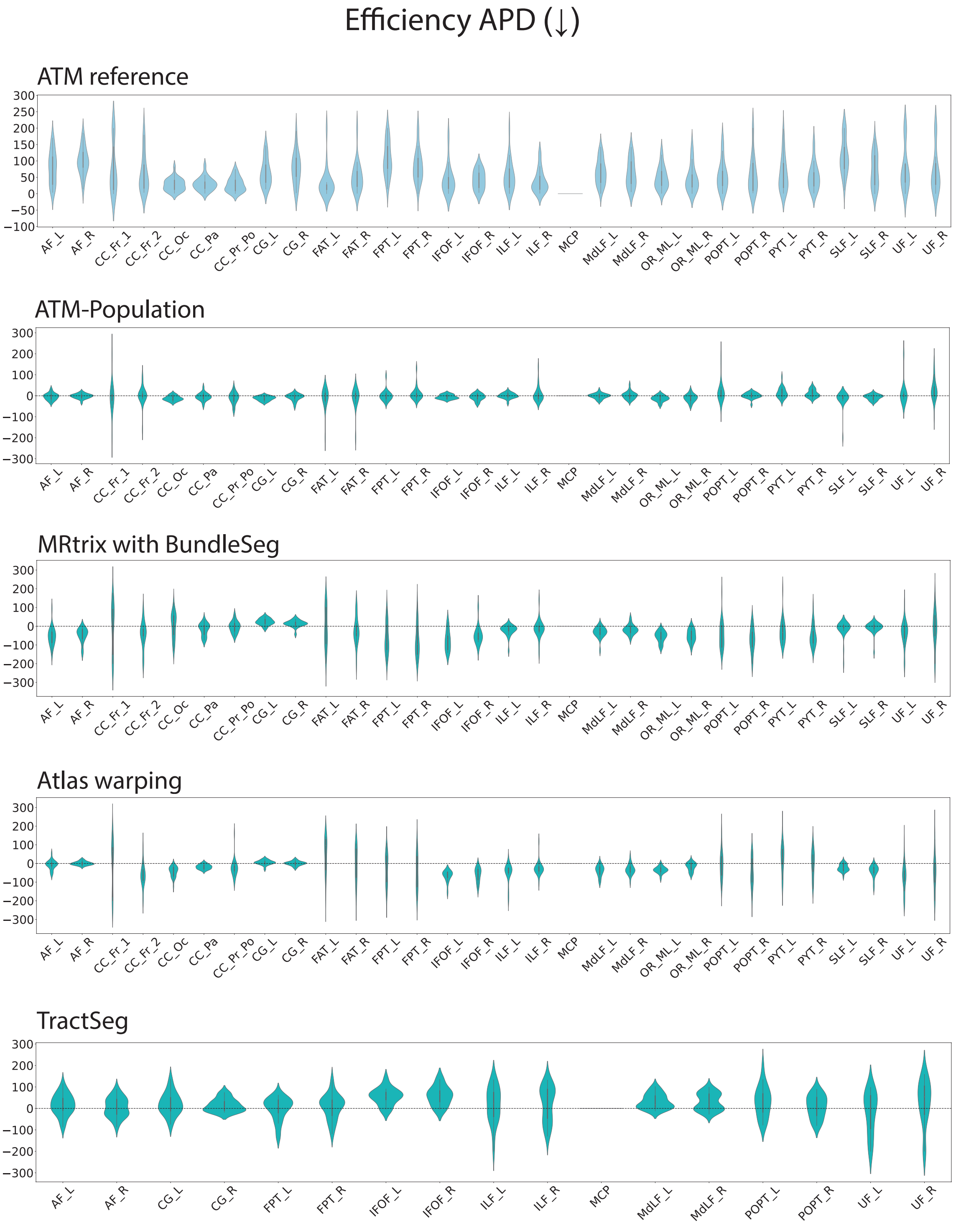


**Fig. S28 | Network global efficiency APD of bundle connectomes generated with ATM, ATM-Population, MRtrix with BundleSeg, SCIL WM atlas warping, and TractSeg.** The first violin plot shows the ATM values as the baseline. The subsequent rows display the paired performance differences of other methods relative to this baseline. For each metric, the arrow indicates preference: ↑ higher is better and ↓ lower is better.

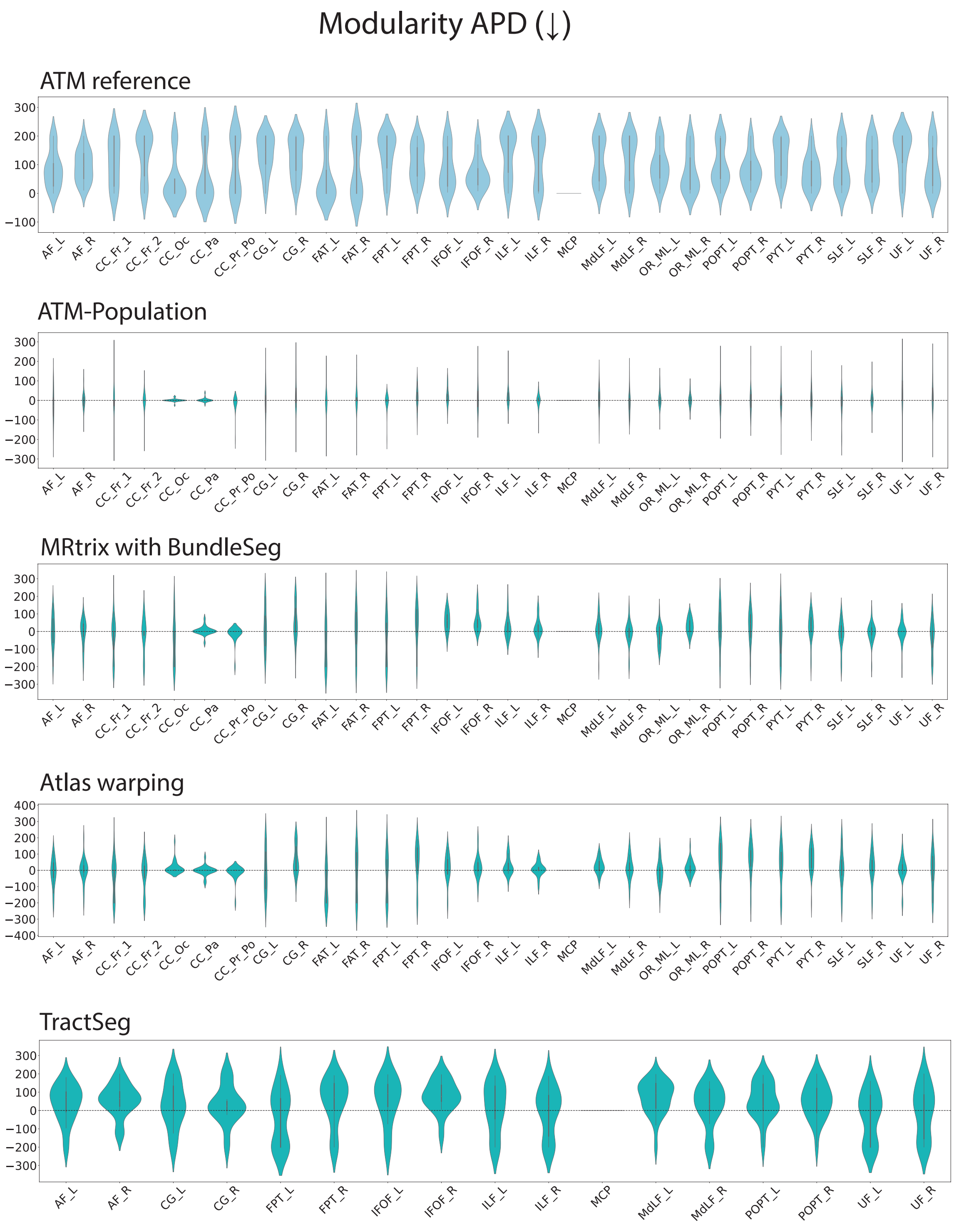


**Fig. S29 | Network modularity APD of bundle connectomes generated with ATM, ATM-Population, MRtrix with BundleSeg, SCIL WM atlas warping, and TractSeg.** The first violin plot shows the ATM values as the baseline. The subsequent rows display the paired performance differences of other methods relative to this baseline. For each metric, the arrow indicates preference: ↑ higher is better and ↓ lower is better.
